# *Anacapa Toolkit*: an environmental DNA toolkit for processing multilocus metabarcode datasets

**DOI:** 10.1101/488627

**Authors:** Emily E. Curd, Zack Gold, Gaurav S Kandlikar, Jesse Gomer, Max Ogden, Taylor O’Connell, Lenore Pipes, Teia Schweizer, Laura Rabichow, Meixi Lin, Baochen Shi, Paul Barber, Nathan Kraft, Robert Wayne, Rachel S. Meyer

## Abstract

1. Environmental DNA (eDNA) metabarcoding is a promising method to monitor species and community diversity that is rapid, affordable, and non-invasive. Longstanding needs of the eDNA community are modular informatics tools, comprehensive and customizable reference databases, flexibility across high-throughput sequencing platforms, fast multilocus metabarcode processing, and accurate taxonomic assignment. As bioinformatics tools continue to improve, addressing each of these demands within a single bioinformatics toolkit is becoming a reality.

2. We present the modular metabarcode sequence toolkit Anacapa (https://github.com/limey-bean/Anacapa/), which addresses the above needs, allowing users to build comprehensive reference databases and assign taxonomy to raw multilocus metabarcode sequence data A novel aspect of Anacapa is our database building module, Creating Reference libraries Using eXisting tools (*CRUX*), which generates comprehensive reference databases for specific user-defined metabarcode loci. The Quality Control and Dereplication module sorts and processes multiple metabarcode loci and processes merged, unmerged and unpaired reads maximizing recovered diversity. Followed by amplicon sequence variants (ASVs) detection using *DADA2*. The Anacapa Classifier module aligns these ASVs to CRUX-generated reference databases using *Bowtie2*. Taxonomy is assigned to ASVs with confidence scores using a Bayesian Lowest Common Ancestor (*BLCA*) method. The *Anacapa Toolkit* also includes an R package, ranacapa, for automated results exploration through standard biodiversity statistical analysis.

3. We performed a series of benchmarking tests to verify that the Anacapa Toolkit generates comprehensive reference databases that capture wide taxonomic diversity and that it can assign high-quality taxonomy to both MiSeq-length and Hi-Seq length sequence data. We demonstrate the value of the *Anacapa Toolkit* to assigning taxonomy to eDNA sequences from seawater samples from southern California including capability of this tool kit to process multilocus metabarcoding data.

4. The *Anacapa Toolkit* broadens the exploration of eDNA and assists in biodiversity assessment and management by generating metabarcode specific databases, processing multilocus data, retaining all read types, and expanding non-traditional eDNA targets. Anacapa software and source code are open and available in a virtual container to ease installation.

## 1 Introduction

Rapid and inexpensive biodiversity monitoring tools are critical for maintaining healthy ecosystems and for effective species conservation (Deiner *et al.* 2017). Environmental DNA (eDNA) is a promising non-invasive approach for biodiversity monitoring in terrestrial and aquatic ecosystems because it provides rapid, cost-effective surveys of a broad array of taxa (Taberlet *et al.* 2012; Deiner *et al.* 2017; Bohmann *et al.* 2014; Kelly *et al.* 2014). eDNA metabarcoding is increasingly used in ecology and conservation research, but three key challenges remain in sequence processing and taxonomic assignment stages, limiting the accuracy and reliability of this approach.

To capture broad taxonomic diversity, many eDNA studies simultaneously sequence multiple loci per sample (e.g. Stat *et al.* 2017). However, few metabarcode pipelines are explicitly designed to process multilocus high-throughput sequencing data (but see Arulandhu *et al.* 2017). To process multilocus eDNA sequence data, researchers must sort and process unique metabarcodes independently, which increases valuable user and computation time for each additional metabarcode.

Another challenge for eDNA metabarcode processing is the lack of robust, locus-specific reference databases (Deiner *et al.* 2017). Curated databases for select metabarcode loci offer validated solutions for certain commonly-used universal metabarcodes (e.g. UNITE Community, 2017), but such curated databases are unlikely to exist for all loci used in metabarcoding studies, especially as the number of target metabarcodes grows. Custom, user-generated databases offer one potential solution, but current approaches can be problematic. For example, generating reference databases through *in silico* PCR will miss reference sequences that do not contain primer recognition sites – which is the case for many sequences available in Genbank (Ficetola *et al.* 2010; Boyer *et al.* 2016). Methods that rely on keyword searches to generate reference databases are sensitive to inaccurate metadata (Machida *et al.* 2017) and are susceptible to retrieving sequences that lack the target metabarcode locus. Together, these issues highlight a need for more comprehensive reference databases that enhance taxonomic assignment for eDNA metabarcoding.

A third challenge of existing metabarcode pipelines is that they frequently discard large portions of sequence data, including reads that can be valuable for assigning taxonomy. For example, many existing pipelines discard unmerged sequences entirely, or only use partial sequence data where full-length alignment with reference metabarcodes is not possible (Port *et al.* 2015). Other tools independently assign taxonomy to unmerged forward and reverse reads assigning the least common ancestor between calls an approach which can lead to poorly-resolved taxonomic assignments (Huson *et al.* 2016). The quality of assignments generated by one pipeline designed to handle unmerged paired data (Bengtsson-Palme *et al.* 2015) suffers from using non-contiguous sequences and suboptimal BLAST parameters (Shah *et al.* 2018). Many pipelines discard significant portions of sequencing data (e.g. remove all reverse reads) for unmerged paired data, potentially causing selection bias against certain taxa (Deagle *et al.* 2014). Although k-mer based classifiers can be modified to accept unmerged paired data, there is no specific documentation or benchmarking for this functionality (e.g. Edgar 2016). Furthermore, the inability of pipelines to incorporate large amounts of sequence information for taxonomic calls will be exacerbated with the use of new high-throughput sequencing platforms (e.g. Illumina NovaSeq and 10X), which generate greater read depth but shorter sequences, which increases the risk of generating unmerged paired data for long metabarcodes.

To help resolve these challenges, we developed the *Anacapa Toolkit*, which contains modules for 1) creating custom reference databases, 2) executing quality control and multilocus read parsing, 3) generating taxonomic assignments for all quality reads produced by HiSeq and MiSeq Illumina platforms, and 4) interactively visualizing taxonomy tables from the *Anacapa Toolkit* using the R package *ranacapa*, described in Kandlikar *et al.* (2018).

## 2 Anacapa Toolkit

The *Anacapa Toolkit* combines components of leading bioinformatics software with custom methods (Figure 1). The first module, Creating Reference libraries Using eXisting tools (*CRUX*), generates custom reference databases. The second module performs quality control of raw sequences and infers Amplicon Sequence Variants (ASVs) using *DADA2* (Callahan *et al.* 2016). The third module assigns taxonomy using *Bowtie2* and the Bayesian Lowest Common Ancestor algorithm (*BLCA*; Gao *et al.* 2017).

**Figure 1.**
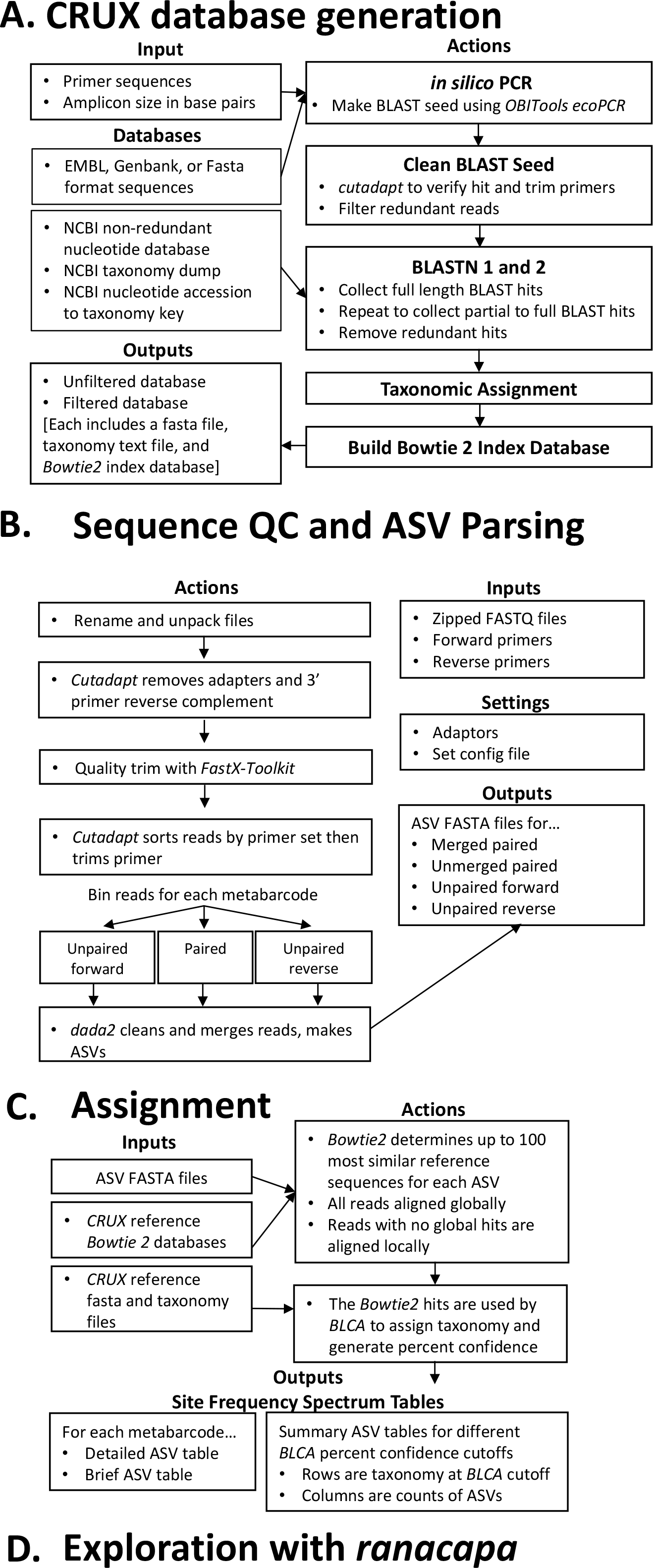
Flowchart of the *Anacapa Toolkit*.

All components of the *Anacapa Toolkit* are openly available (https://github.com/limey-bean/Anacapa; DRYAD link provided upon acceptance). The *Anacapa Toolkit* and several *CRUX*-generated reference databases are available in virtual containers developed with Code for Science and Society (Ogden 2018).

### 2.1 *CRUX*: Creating Reference libraries Using eXisting tools

The *Anacapa Toolkit’s* first module, *CRUX* (Figure 1A; Appendix 1.1), constructs custom reference databases for user-defined primers by querying public databases. *CRUX* first generates metabarcode-specific seed databases by running *in silico* PCR (Ficetola *et al.* 2010) on the EMBL standard nucleotide database (Stoesser *et al.* 2002). To increase the breadth of reference sequences and capture sequences without barcode primers, *CRUX* then uses BLASTn (Camacho *et al.* 2009) to query the seed databases against the NCBI non-redundant nucleotide database (Pruitt *et al.* 2005). *CRUX* de-replicates the BLASTn hits by retaining only the longest version of each sequence and retrieves taxonomy using Entrez-qiime (Baker, 2016). For each primer set, *CRUX* generates an “unfiltered” database which contains all accessions and taxonomic paths, a paired “filtered” database which excludes accessions with ambiguities in the taxonomic paths, and a *Bowtie2-formatted* index library (Langmead & Salzberg, 2012).

### 2.2 Sequence quality control and ASV parsing

The *Anacapa Toolkit’s* Quality Control and De-replication module (Figure 1B; Appendix 1.2) conducts standard DNA sequence quality control and generates ASVs. It uses *cutadapt* (Martin, 2011) and *FastX-toolkit* (Gordon & Hannon, 2010) to trim user-defined primers and adapters and low-quality bases from raw FASTQ files from Illumina sequencing platforms. Next, this module uses *cutadapt* to separate reads from multiple loci within each sample. A custom Python script sorts locus-specific reads into three categories: paired-end reads, forward-only reads, and reverse-only reads. These reads are then processed separately through *DADA2* (Callahan *et al.* 2016) to denoise, dereplicate, merge paired reads, and remove chimeric sequences. This step returns ASV FASTA files and ASV count summary tables for four read types: merged paired-end reads, unmerged paired-end reads (filtered based on length and overlap criteria; Appendix 1.2), forward-only reads, and reverse-only reads. These files are inputs for the *Anacapa Toolkit* taxonomic assignment module.

### 2.3 Taxonomic assignment: *Anacapa Classifier* assigns taxonomy with *Bowtie2* and *BLCA*

The Anacapa Classifier module (Figure 1C; Appendix 1.3) assigns taxonomy to ASVs using *Bowtie2* and a modified version of *BLCA* (Gao *et al.* 2017). We verified that our modification to *BLCA* (namely, accepting Bowtie2-formatted SAM files rather than BLAST output files) does not influence taxonomy assignment (Appendix 4). In the first step of the Anacapa Classifier, *Bowtie2* queries ASVs against metabarcode-specific *CRUX* generated reference databases returning up to 100 alignments per ASV. The module uses *Bowtie2’s* “very-sensitive” preset to ensure high-quality alignments. The *Bowtie2* outputs are then processed with *Bowtie2-BLCA*, using multiple sequence alignment to probabilistically determine taxonomic identity by selecting the lowest common ancestor from the multiple weighted *Bowtie2* hits for each ASV. This module returns both detailed and brief reports of taxonomic assignment, and eight sets of taxonomy tables based on varying bootstrap confidence cutoffs (40-100).

## 3 Benchmarking the *Anacapa Toolkit*

To benchmark the performance of the first three modules of the *Anacapa Toolkit*, we performed a series of quantitative tests of these modules on various metabarcodes and sequencing read types. We found that *CRUX*-generated databases capture greater taxonomic diversity than published marker-specific reference databases for *CO1*, *12S*, and Fungal *ITS* metabarcodes (Appendix 3). We verified that the Anacapa Classifier consistently generates high-quality taxonomic assignments and explored the consequence of varying bootstrap confidence cutoff on assigned taxonomy (Appendix 4). Finally, we verified that the *Anacapa Toolkit* Quality Control and Classifier modules can process both longer (e.g. MiSeq) and shorter (e.g. HiSeq) DNA metabarcoding sequences (Appendix 5).

## 4 Case Study: Using the *Anacapa Toolkit* to assign taxonomy to field-collected eDNA samples

To test the *Anacapa Toolkit* on field-collected eDNA metabarcoding datasets, we processed 30 seawater samples from kelp forests across the southern California Channel Islands. Seawater samples were amplified using 12S (Miya *et al.* 2015) and CO1 (Leray *et al.* 2013) metabarcodes (see Appendix 6 for laboratory preparation and data analysis; Table S6.3). For the 12S metabarcode, the *CRUX* module was critical for assigning taxonomy as there are no published reference databases that included all vertebrate species for this locus (Sato *et al.* 2018). Sequence data from these samples are available in NCBI (SRA accession SRP140860). Across seawater samples, we generated 15,745,317 paired-end sequencing reads, of which 11,866,904 reads and 37,910 ASV were identified as 12S and 3,878,413 reads and 15,997 ASVs were identified as CO1. For both metabarcodes, we found that 99.5% were merged read pairs and <1% were unmerged paired, forward or reverse only reads. The *Anacapa Toolkits* taxonomic assignments included 533 assignments at the species level, 414 Genera, 295 Families, 49 Classes, and 21 Phyla of eukaryotes. This included many taxa of interest for natural resource managers (Figure 2; Tables S6.1,S6.2), highlighting the ability of eDNA to detect a wide breadth of marine life and its utility for biodiversity monitoring. A detailed and interactive summary of these seawater samples is available as the demo dataset of the *ranacapa* module (https://gauravsk.shinyapps.io/ranacapa/).

**Figure 2.**
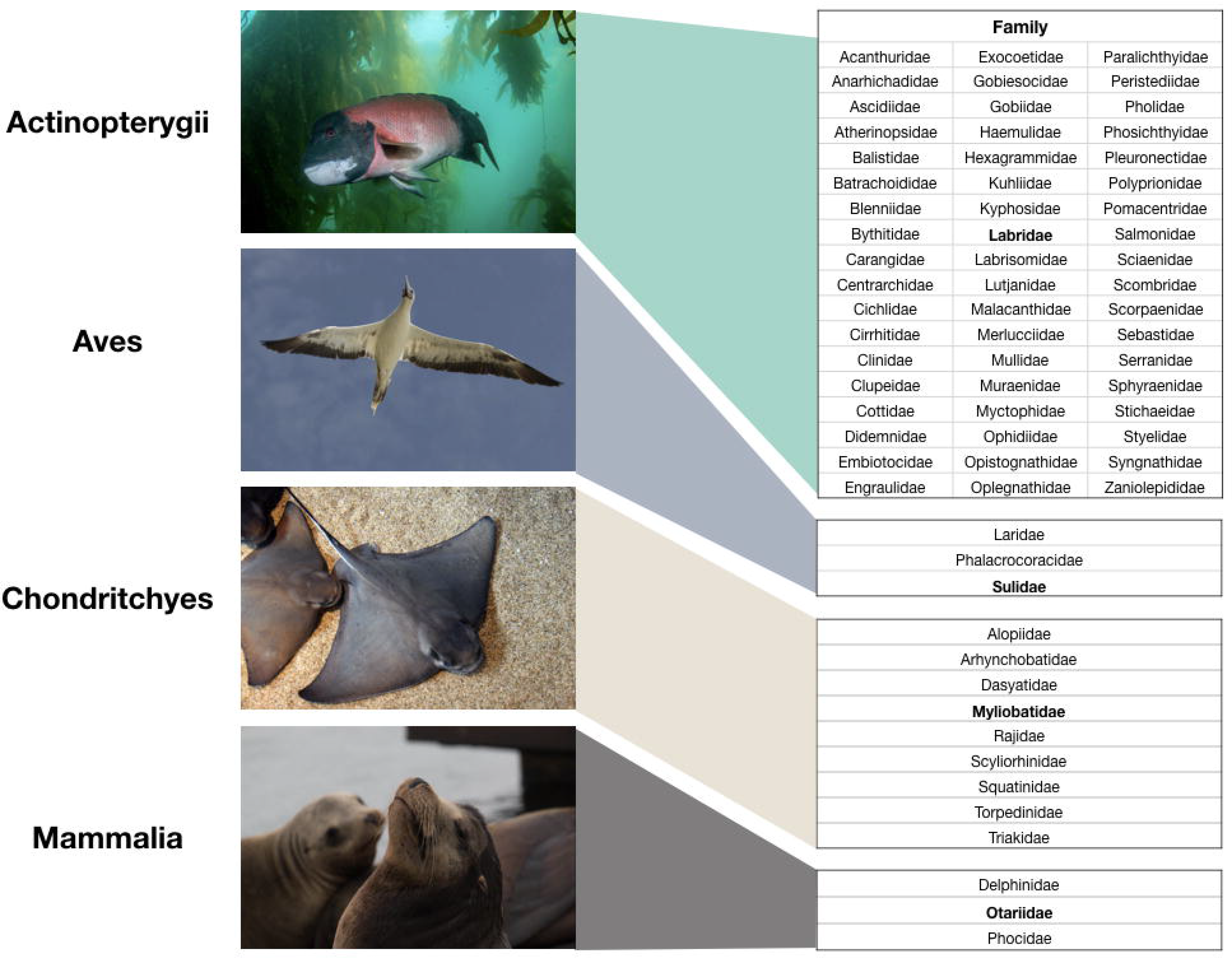
Taxonomic assignments from California environmental samples. **A.** Highlights 12S metabarcodes of Anacapa Island kelp forest vertebrate families. Families in bold are featured.

## 5 Conclusion

Biodiversity monitoring initiatives are increasingly using eDNA to inventory communities using multilocus metabarcoding. However, a bottleneck in the broad use of eDNA approaches is the lack of bioinformatic pipelines that are accurate and easily customizable. The *Anacapa Toolkit* provides such functionality for eDNA projects as well as other common applications such as gut content analysis (Leray *et al.* 2013) and autonomous reef monitoring structures (Ransome *et al.* 2017). The *Anacapa Toolkit* is modular and its parameters are easily modifiable, so that it can be adapted to users’ specific needs in several important ways. First, *CRUX* reference databases are compatible with alternative classifiers (Bokulich *et al.* 2018), and users can append their own reference sequences to *CRUX* databases as needed. Second, the quality control and ASV parsing module are designed to process pooled metabarcoding libraries and automatically sort them by barcode and sample. The resulting output files (with the exception of unmerged reads) can be analyzed by most classifiers. Third, the *Anacapa Bowtie2-BLCA* Classifier can be applied to any high-throughput sequencing data, and process paired and unpaired reads. The robustness of CRUX-generated reference databases and the flexibility of the *Anacapa Toolkit* enables eDNA studies with a variety of metabarcodes to efficiently assign taxonomy in a wide range of community ecology and biodiversity management research.

## Supporting information

## Acknowledgements

The *Anacapa Toolkit* is a product of the CALeDNA community science program (www.ucedna.com). We thank R. Turba, D.R. Ramos, E.W. Rankin, S. Stinson, and S. Shirazi for beta-testing, B. Shapiro for advice, and J.S. Guswa for editing the manuscript. We thank R. D’Auria for assistance in implementing Anacapa on the UCLA cluster, and S. Chadalapaka for assistance at UC-Merced. We thank D. Kushner and J. Sprague for sample collection. This work was funded by the University of California President’s Research Catalyst Award (CA-16-376437), NSF-DEB 1644641 (NK), NSF-DGE 1650604 (GK), NSF-GRFP 2015204395 (ZG). The Gordon and Betty Moore Foundation (6864) supported the Dat in the Lab project that containerized Anacapa.

## Author Contributions

EEC coordinated toolkit development and benchmarking. BS, EEC, GSK, JG, RSM, RW, and ZG designed Anacapa. BS, EEC, GK, JG, MO, and ZG wrote Anacapa. EEC and MO built the Singularity container. EEC, GK, JG, LP, ML, RSM, RW, TO, and ZG designed or conducted benchmarking studies. EEC, LR, TS, and ZG generated the California eDNA libraries. EEC, GK, and ZG wrote the manuscript with help from LP, LR, ML, RSM and TO; PB, RW, TS revised. NK, PB, and RW provided resources to support the work.

