## Supplementary material for "*Anacapa Toolkit*: an environmental DNA toolkit for processing multilocus metabarcode datasets"

**Appendix 1. *Anacapa Toolkit* parameters**

*1.1 Generating CRUX reference databases*

*CRUX* databases were built using the *CRUX* default parameters described below (also see <https://github.com/limey-bean/CRUX_Creating-Reference-libraries-Using-eXisting-tools>; version will be permanently archived in Dryad pending acceptance). All user-modifiable parameters are underlined. Loci and primer pairs used to generate reference databases are found in Table S2.1. All *CRUX*-generated reference databases can be downloaded at (Dryad link will be provided pending acceptance).

The first step in *CRUX* is to generate an *OBItools* (Boyer et al. 2016) compatible EMBL library (ftp.ebi.ac.uk/pub/databases/embl/release) that will be used to generate a seed library of matching barcode targets. To do this we downloaded EMBL ENA Release 133 sequence collections that represent a broad diversity of life: Invertebrates (INV), plants (PLN), prokaryotes (PRO), fungi (FUN), and other vertebrates (VRT). Seed libraries were built using *ecoPCR* (Ficetola et al. 2010). For seed libraries constructed with primers without degenerate base pairs, we allowed 3 mismatches between primers and reference. This accounts for expected primer mismatch using universal primer sets (Deagle et al. 2014). However, *ecoPCR* does not take degenerate bases into account, and thus for primers with degenerate bases we increased the number of mismatches allowed to equal the maximum number of degenerate bases present either of the primers. For example, the reverse CO1 primer (jgHCO2198) has 8 degenerate sites and the forward CO1 (mlCOIintF) has 7 degenerate sites (Table 2.1), therefore the number of allowed mismatches for CO1 was 8.

For *ecoPCR*, we set the minimum expected amplicon length and maximum amplicon length as 100 base pairs fewer or greater than the average length published in previous reports (see references in Table S1.1). We then verified that the both primers were present in each sequence, and subsequently trimmed off the primers using *cutadapt* (version 1.16; Martin, 2011) allowing for 30% mismatch between primers and query. Given that *cutadapt* takes into account sequence degeneracy, this threshold was held constant for each primer set.

We then used these cleaned sequences as a seed set for blastn 2.6.0+ (Camacho et al., 2009). Nucleotide (nt) BLAST databases (ftp.ncbi.nlm.nih.gov/blast/db/) were downloaded on November 08, 2017. *blastn* was run twice on the seed sets, using these default settings (minimum e-value 0.00001, 10 threads, maximum return of 10,000 matches). The first *blastn* run retrieved matches with a minimum 50% identity and were required to match the entire length of the subject. The second run retrieved matches with a minimum 70% identity and minimum of 70% coverage of the subject, therefore retaining many partial sequences. Returned sequences were then concatenated, de-replicated so as to keep only the longest read for a unique NCBI accession version identifier. Sequences were converted to FASTA format and a taxonomy file was generated based on the accession version using *entrez_qiime.py* (Baker, 2016). The fasta file and corresponding taxonomy file were deemed the ‘unfiltered *CRUX* database’. We then made a ‘filtered *CRUX* database’ by removing FASTA sequences with ambiguous taxonomic assignments (i.e. taxonomic paths that include the phrase “uncultured”, “environmental”, “sample”, or four consecutive missing identifications, i.e. “NA;NA;NA;NA”). Databases were indexed with *Bowtie 2* formatting using bowtie2-build (Langmead et al., 2012).

The filtered *CRUX*-generated *16S, 18S, CO1, PITS, FITS*, and *12S* libraries from Table S2.1 are available at <https://github.com/datproject/anacapa-container> and (dryad link available upon publication).

*1.2 Anacapa Sequence QC and ASV Parsing*

Raw FASTQ sequence libraries are run through the *Anacapa Sequence QC and ASV Parsing* step of the *Anacapa Toolkit* using the default parameters described below. All user modifiable parameters are underlined. Here we describe the processing steps for both types of read inputs: HiSeq and MiSeq.

The following steps are automated in the running of the *Anacapa Sequence QC and ASV parsing Software* (see anacapa_QC_dada2.sh script in https://github.com/limey-bean/Anacapa). Raw Illumina reads (.fastq.gz) are given a digital fingerprint (*md5sum*), uncompressed, and the file names are shortened by replacing the standard R1_001.fastq and R2_001.fastq suffixes to _1.fastq and _2.fastq. If not using the default primer sets (*12S, 18S, 16S, FITS, PITS,* and *CO1*), paths to primer sequence(s) and expected merge length files need to be entered by the user in the command line fields. Adapter type (Nextera, TruSeq, or NEBnex) is also indicated by the user in command line fields. The default settings used to process data are as follows: We allow up to 30% error in match between sequence and adapter or primer(s) + sequence adapters. Then *cutadapt* removs the 5’ sequence adapter and 3’ primer(s) + sequence adapter allowing for a 30% error rate. The 3’ primer(s) + sequence adapter is removed because the metabarcode read may be shorter in length than the sequencing platform read length, producing sequences that extend through the end of the 3’ primer and sequence adapter. We remove low quality sequence (quality score < 35) using fastq_quality_trimmer (*FastX_toolkit* Version: 0.0.13; Gordon and Hannon, 2010). Next, reads are sorted into metabarcode specific directories based on the 5’ primer sequence(s), and only reads > 99 bp are retained.

We then perform an additional trimming step for all reads to improve read merging in *DADA2* (see below; Callahan et al. 2016). The *DADA2* documentation recommends that users trim some length of base pairs from the end of reads based on read quality profile. Therefore, we trim an additional 20 bp from MiSeq forward reads, 50 bp from MiSeq reverse reads, 10 bp from HiSeq forward reads and 25 bp from HiSeq reverse reads. We remove more bases from reverse reads as reverse Illumina reads tend to be of lower quality (<https://www.illumina.com/documents/products/technotes/technote_Q-Scores.pdf>).

We then implement a custom *Python 2-4.2* (<http://biopython.org/wiki/Packages>*)* script to sort reads in metabarcode directories into unpaired forward, unpaired reverse, and paired read directories.

These sorted reads are then de-replicated into ASVs using *DADA2*, following the steps of the *DADA2* tutorial for paired end reads (<https://benjjneb.github.io/dada2/tutorial.html>), but with a few exceptions and modifications. First, given we have pre-trimmed reads, we do not trim reads in the “filterAndTrim” step. Second, we modified the script to accept unmerged paired reads. Typically unmerged paired end reads are discarded in *DADA2*. However, to maximize data retention we keep unmerged paired end reads as long as the total length of both read pairs was less than the expected length of the amplicon (without primers) plus 20 base pairs. This is the expected minimum length for paired reads to overlap in *DADA2* for read merging. We then retain the unmerged reads that passed the length criteria, and check them for chimeras and remove chimeras using the following method. To allow unmerged paired reads to be processed for chimera detection, a dummy sequence (“AAAAAAAAAATTCTTAAAAAAAAAA”) is added between the forward and reverse complement of the reverse read. Modified reads (unmerged F + dummy sequence + reverse complement unmerged R) are then run through the *DADA2* chimera checker. The dummy sequence is removed from all non-chimeric unmerged reads and the remaining reads are binned in two FASTA files: unmerged forward and unmerged reverse reads. We also retain *DADA2* processed unpaired forward and reverse reads. For unpaired forward and reverse reads, we run all of the *DADA2* steps in the same order as for the merged reads but omit the read merging and retention of unmerged reads steps.

The final results of *DADA2* are tables of unique ASVs with sequences and counts of ASV per sample for each read type (unpaired forward, unpaired reverse, merged, unmerged). FASTA files of ASV reads are also given; one file per unpaired forward, unpaired reverse read types, and two files for unmerged read types (unmerged forward and reverse). Example datasets and outputs are available at <https://github.com/datproject/anacapa-container> and <https://github.com/limey-bean/Anacapa>.

*1.3 Anacapa Bowtie2-BLCA Classifier*

ASVs generated in the *Anacapa Sequence QC and ASV Parsing* step were run through the *Anacapa Bowtie2-BLCA Classifier* using the default parameters. All user-modifiable parameters are underlined below.

The following steps are automated in the running of the *Anacapa Bowtie2-BLCA Classifier* Software (see anacapa_classifier.sh script in https://github.com/limey-bean/Anacapa). The first step in taxonomic assignment is to map the ASVs generated in in the *Anacapa Sequence QC and ASV Parsing* step to *CRUX* references using *Bowtie2*. Reads are first mapped using *Bowtie2*’s global alignment mode “end-to-end”. Any reads that do not map in end-to-end mode are then mapped using local alignment mode. *Bowtie2*’s “very-sensitive” setting is used in both modes, and other parameters included setting 120 threads, and retention of a maximum of 100 matches per query. The use of *Bowtie2’s* “very-sensitive” parameter produces the highest possible alignment accuracy for the method. For unmerged reads, we used an additional parameter that forced *Bowtie2* to only consider alignments that contained both read pairs.

The *Bowtie2* matches are then passed into the Bayesian Least Common Ancestor (*BLCA*) classifier (Gao et al. 2017). In brief, the classifier generates multiple sequence alignments between the query and database matches. Each database match contributes a weighted Bayesian posterior probability for a taxonomic assignment, where dissimilar matches are penalized. Taxonomy is ultimately assigned based on the lowest common ancestor of multiple weighted reference library matches for each query sequence. The reliability of each taxonomic assignment is then evaluated through bootstrap confidence scores. The default level of Bootstrapping is 100 random samplings with replacement. Only reads with a minimum percent identity (-b 80%) and a minimum percent subject length relative to the query (-p 80%) were passed into BLCA. The resulting files for the *Anacapa Bowtie2-BLCA classifier* are summary tables that concatenate the classification results from forward, reverse, merged and unmerged paired reads according to their assigned taxonomy, and summary tables with taxonomic rank reported if the bootstrap confidence cutoff is above a certain value (e.g 100, 95, 90, 80, 70, 60, 50, and 40).

Example Datasets and outputs are available at <https://github.com/datproject/anacapa-container> and <https://github.com/limey-bean/Anacapa>.

*1.4 Running ranacapa*

To begin exploring this output, we wrote the R package *ranacapa (*Kandlikar et al. 2018), which relies heavily on the packages *phyloseq* (McMurdie and Holmes 2013) and *vegan* (Oksanen *et al.* 2017) to rarefy, visualize, and conduct exploratory statistics. The package is available for installation at <https://github.com/gauravsk/ranacapa> using the command devtools::install_github("gauravsk/ranacapa"). *ranacapa* utilities can be run either as an interactive *Shiny* webapp (https://gauravsk.shinyapps.io/ranacapa/) or through the command line using the *Anacapa* script ranacapa_automated.R.
