## Supplementary material for "*Anacapa Toolkit*: an environmental DNA toolkit for processing multilocus metabarcode datasets"

**Appendix 2. Metabarcodes and datasets**

*2.1 Metabarcode primer sets*

In this study we benchmarked the *Anacapa Toolkit* using 8 metabarcode loci; *12S* MiFish (Miya *et al.* 2015), *16S* V4 (Caporaso *et al*., 2012), *18S* V4 (Stoeck *et al*. 2010; Bradley *et al*. 2016), *18S* V8-9 (Bradley *et al*. 2016), *18S* (Amaral-Zettler *et al*. 2009), Plant ITS2 “*PITS*” (Gu *et al*. 2013), *CO1* (Leray *et al*. 2013, Geller et al. 2013), and Fungal ITS “*FITS*” (White *et al.* 1990; Epp *et al.* 2012). See Table S2.1 for primer details.

*2.2. Isolate sanger datasets used in reference database and classifier benchmarking*

We used several published isolate datasets from mock community analyses to benchmark the *Anacapa toolkit* (Table S2.2). For database and classifier comparisons, we used Sanger generated reads from isolates for *16S* (n=75 unique isolates: Krohn *et al.* 2016, Callahan *et al.* 2016, Kozich *et al.* 2013), *CO1* (n=34 organisms: Leray *et al.* 2017), and two *18S* (V4 n=12 and V8-9 n=12 organisms: Bradley *et al.* 2016) metabarcode loci.

*2.3. Mock community datasets used in MiSeq and pseudo-HiSeq benchmarking*

For taxonomic assignment comparison between MiSeq and pseudo-HiSeq datasets, we selected mock communities generated from isolate sequence pools for *16S* (Callahan *et al.* 2016), *CO1* (Leray *et al.* 2017), and two *18S* (Bradley *et al.* 2016) loci that were run on a MiSeq platform. Isolates represented in the Callahan *et al.* (2016) *16S* mock community (n = 35 isolates, comprised of 27 species), were pooled at different orders of magnitude and the exact ratio can be found in Table S5.6. Isolates represented in the Leray *et al.* 2017 *CO1* mock community (n = 34 isolates comprised of unique species) consist of the entire “run 1” mock data set in which all 34 isolates were evenly pooled (see Table S5.7). Isolates represented in the Bradley *et al*. 2016 *18S* V4 and V8-9 mock communities (total n = 12 species) were pooled at different concentrations (see Tables S5.8 and S5.9), and we selected four mock communities to analyze in the current study; MC11 (n=6 organisms), MC21 (n=12 organisms), MC51 (n=12 organisms), and MC61 (n=12 organisms).

*2.4. Field collected eDNA datasets used in the case study*

Environmental DNA datasets generated from CALeDNA (ucedna.com) seawater samples were used to demonstrate the *Anacapa Toolkit* with field collected eDNA data from CALeDNA, to benchmark processing time for the *Anacapa Bowtie2-BLCA* and the original *BLAST-BLCA* classifier. In total, 30 samples were collected (Table S6.3) around the Southern California Channel Islands including Anacapa Island. The raw sequencing data is available in the NCBI SRA (SRP140860).

throughput microbial community analysis on the Illumina HiSeq and MiSeq

platforms. *The ISME Journal*, **6**,1621–1624. doi: 10.1038/ismej.2012.8 PMID:

22402401.

Epp, L.S., Boessenkool, S., Bellemain, E.P., Haile, J., Esposito, A., Riaz, T., Erseus, C., Gusarov, V.I., Edwards, M.E., Johnsen, A. and Stenøien, H.K. (2012). New environmental metabarcodes for analysing soil DNA: potential for studying past and present ecosystems. *Molecular Ecology*, **21**, 1821-1833.

Geller, J., Meyer, C., Parker, M. and Hawk, H. (2013). Redesign of PCR primers for mitochondrial cytochrome c oxidase subunit I for marine invertebrates and application in all‐taxa biotic surveys. *Molecular Ecology Resources*, **13**, 851-861.

Gu, W., Song, J., Cao, Y., Sun, Q., Yao, H., Wu, Q., Chao, J., Zhou, J., Xue, W. and Duan, J. (2013). Application of the ITS2 region for barcoding medicinal plants of Selaginellaceae in Pteridophyta. *PloS One*, **8**, e67818.

Kozich, J.J., Westcott, S.L., Baxter, N.T., Highlander, S.K. and Schloss, P.D. (2013). Development of a dual-index sequencing strategy and curation pipeline for analyzing amplicon sequence data on the MiSeq Illumina sequencing platform. *Applied and Environmental Microbiology*, **79**, 5112-5120.

Krohn, A., Stevens, B., Robbins-Pianka, A., Belus, M., Allan, G.J. and Gehring, C. (2016). Optimization of 16S amplicon analysis using mock communities: implications for estimating community diversity. *PeerJ Preprints*, **10**.

Leray, M., Yang, J.Y., Meyer, C.P., Mills, S.C., Agudelo, N., Ranwez, V., Boehm, J.T. and Machida, R.J. (2013). A new versatile primer set targeting a short fragment of the mitochondrial COI region for metabarcoding metazoan diversity: application for characterizing coral reef fish gut contents. *Frontiers in Zoology*, **10**, 34.

Leray, M., & Knowlton, N. (2017). Random sampling causes the low reproducibility of rare eukaryotic OTUs in Illumina COI metabarcoding. *PeerJ*, **5**, e3006. doi:10.7717/peerj.3006

Miya, M., Sato, Y., Fukunaga, T., Sado, T., Poulsen, J.Y., Sato, K., Minamoto, T., Yamamoto, S., Yamanaka, H., Araki, H. and Kondoh, M. (2015). MiFish, a set of universal PCR primers for metabarcoding environmental DNA from fishes: detection of more than 230 subtropical marine species. *Royal Society Open Science*, **2**, 150088.

Stoeck, T., Bass, D., Nebel, M., Christen, R., Jones, M.D., Breiner, H.W. and Richards, T.A. (2010). Multiple marker parallel tag environmental DNA sequencing reveals a highly complex eukaryotic community in marine anoxic water. *Molecular Ecology*, **19**, 21-31.

White, T.J., Bruns, T., Lee, S.J.W.T. and Taylor, J.L. (1990). Amplification and direct sequencing of fungal ribosomal RNA genes for phylogenetics. *PCR protocols: a guide to methods and applications*, **18**, 315-322.
