## Supplementary material for "*Anacapa Toolkit*: an environmental DNA toolkit for processing multilocus metabarcode datasets"

**Appendix 3. Database Performance of *CRUX* and Published Reference Databases**

*3.1 Database performance of CRUX and published reference databases*

We benchmarked metabarcode-specific reference databases generated by *CRUX* by comparing these databases to published reference databases for corresponding metabarcodes. We used default *CRUX* parameters to generate reference libraries for three commonly-used eDNA metabarcodes: the CO1 metabarcode (Leray et al. 2013; hereafter “*CRUX*-CO1”), the 12S 155 MiFish Universal Teleost metabarcode (Miya et al. 2015; hereafter “*CRUX*-12S”), and the Fungal ITS barcode (Epp et al. 2012; hereafter “*CRUX*-FITS”). In each case, filtered versions of *CRUX*-generated reference databases contained more sequences for more species than corresponding published databases regardless of whether or not published databases had been “restricted” to include only reads that contain the metabarcode sequence (Table S3.1; see Appendix 3.3 for details on how published databases were restricted). We also found that published databases in our comparison have longer average sequence length than corresponding *CRUX* generated databases (Table S3.1 & Figure S3.1).

To further test the quality of *CRUX*-generated reference libraries, we performed a series of pairwise comparisons of taxonomic and phylogenetic breadth, and classification ability between *CRUX*-generated and either unrestricted and restricted published databases (Sections 3.2.1-3.2.3; Appendix 3.2). We performed additional benchmarking with 16S and 18S metabarcode databases and tested taxonomic assignment of Sanger sequenced isolates using 16S, 18S, and CO1 reference databases (Appendices 2.2 and 3.6).

3.1.1 | *CRUX*-generated CO1 reference database

We compared sequences contained in *CRUX*-CO1 to three recently published CO1 databases: Midori (Machida et al. 2017); the CO1 database built by Porter and Hajibabaei (2018; hereafter, P-H database); and CO-ARBitrator (Heller et al. 2018; hereafter, CO-ARBitrator database). For these comparisons, we first pruned published databases to retain only reads that included the Leray et al. (2013) region at the CO1 locus (Appendix 3.3). This reduced the number of species in published databases by 20.3%-48.5% (Table S3.1), and allowed for direct comparisons with *CRUX*-CO1 by retaining only reference sequences that align to the CO1 metabarcoding region generated by Leray et al. primers. Hereafter we refer to restricted databases as “R-[databases]” (e.g. R-Midori). *CRUX*-CO1 contained references for almost four times as many species as the R-P-H database, three times as many as R-Midori, and nearly twice as many as the R-CO-ARBitrator database (Table S3.1). To further compare phylogenetic coverage of *CRUX*-CO1 and published CO1 databases, we mapped taxa present in *CRUX*-CO1 and the restricted published databases onto a large phylogeny of Coleoptera, a taxonomically diverse clade (McKenna et al. 2015; Appendix 3.4). We found that *CRUX*-CO1 contained reference sequences from 66% of Coleopteran genera, whereas the three other restricted databases captured between 41-52% of the genera (Figure S3.2; Table S3.2).

To further evaluate the taxonomic breadth of *CRUX*-CO1 relative to the published databases, we used the Anacapa Classifier module to perform reciprocal pairwise database 190 comparisons. For these tests, we used the Anacapa Classifier to assign taxonomy to sequences in *CRUX*-CO1 using restricted published reference databases as the reference, and vice-versa 192 (Appendix 3.5). Nearly all reads in *CRUX*-CO1 aligned to reads in R-Midori, R-P-H, and R-COARBitrator; similarly, nearly all reads in R-Midori and the R-P-H database aligned to reads in *CRUX*-CO1. However, more than 113,000 reads in the R-CO-ARBitrator database did not align to sequences in *CRUX*-CO1 (Table S3.3). Subsequent BLAST searches of unaligned R-COARBitrator CO1 sequences matched off-target hits (e.g. rbcL, matK, rRNA and ITS genes; Appendix 3.5.2).

To further investigate these unaligned sequences, we used HMMER (Finn et al. 2011; Appendix 3.5.3) to build a profile hidden Markov model of 163,325 CO1 sequences downloaded from the Barcode of Life Data System (BOLD; release 6.50; www.boldsystems.org). We then searched the hidden Markov model against all unaligned CO1 reads from the reciprocal database test and found that nearly all unaligned R-CO-ARBitrator sequences were unlikely to originate from the CO1 gene region (Table S3.3). Conversely, we found that the vast majority of *CRUX*-CO1 reads that did not align to published databases were likely true CO1 sequences (Table S3.3).

Finally, we assessed the quality of taxonomic assignments generated by the Anacapa Classifier module during the reciprocal database test on the *CRUX*-CO1 database versus each restricted published CO1 database using metrics implemented in Taxonomic Cross Validation by Identity (TAXXI; Edgar 2018; Appendix 3.5.4). Specifically, we calculated the True Positive Rate (TPR), which summarizes the rate at which correct taxonomy was assigned at each taxonomic rank out of the total number of opportunities for correct classification, and the Accuracy (ACC) metric, which reflects true positive rates and the rate at which classification results in incorrect taxonomic calls. *CRUX*-CO1 had markedly higher TPR and ACC scores than calls made using all restricted published databases at family, genus, and species ranks (average TPR using *CRUX*-CO1 = 89.7; average TPR using published databases = 40.16, Table S3.3, S3.7). These divergent TPR and ACC scores reflect the fact that *CRUX*-CO1 contains references for nearly all species in the published databases, and that conversely, these published databases contain references for fewer species than *CRUX*-CO1.

3.1.2 | *CRUX*-generated 12S reference database

We compared *CRUX*-12S to reference sequences in the MitoFish reference database (Sato et al. 2018; hereafter “MitoFish”). Restricting MitoFish to reads that included the Miya et al. (2015) 12S region reduced species representation by 73.8% (Table S3.1). *CRUX*-12S database contains sequences for almost twice as many overall species as R-MitoFish (Table S3.1). The unique species in the *CRUX*-12S are comprised of 50% bony fishes (Actinopterygii), 2.4 % cartilaginous fishes (Chondrichthyes), 47% other vertebrate taxa, and fewer than 40 invertebrate taxon reads.

To compare phylogenetic coverage of bony fishes by *CRUX*-12S and the R-MitoFish database, we mapped each reference database onto a species-level phylogeny of Teleostei 230 (Betancur-R et al. 2013; Appendix 3.4). Despite differences in overall database size, *CRUX*-12S and the R-MitoFish databases both cover a majority of the teleost phylogeny (Figure S3.2), and for these Teleostei coverage differed by only 29 species. *CRUX*-12S includes 14 teleost species that are absent from the R-MitoFish database, and R-MitoFish includes 15 teleost species that are absent from *CRUX*-12S (Table S3.4).

In reciprocal database comparisons, we found that all sequences in the R-MitoFish database aligned to sequences in *CRUX*-12S. In contrast, over half of reads in the larger *CRUX* 12S database did not align to sequences in the R-MitoFish database, indicating a greater taxonomic coverage (Table S3.3). To confirm whether unaligned sequences were truly 12S sequences, we analyzed all unaligned sequences in barrnap Version 0.9 (http://www.vicbioinformatics.com/software.barrnap.shtml), which predicts the location of rRNA genes in sequences. Results showed that >91% of *CRUX*-12S reads that did not align to the R-MitoFish database were predicted to be 12S genes, with most originating from non-teleost vertebrate taxa. BLASTn identified 81% of these reads as 12S and the other 19% as mitochondrial in origin, suggesting that not all 12S metabarcode reads are predicted by barrnap. Both *CRUX*-12S and the R-MitoFish had very similar TPRs and ACC scores at the family, 246 genus, and species levels (TPR using *CRUX*-12S = 91.1; TPR using MitoFish = 90.7, Table S3.3, S3.5), indicating that for only teleost fishes, both reference databases perform similarly. Taxonomic assignments were improved by including additional vertebrate 12S sequences in the *CRUX*-12S reference database (see Case Study 4.1).

3.1.3 | *CRUX*-generated Fungal ITS reference database

We compared sequences included in *CRUX*-FITS to reference sequences in two published databases for this region: UNITE (UNITE Community, 2017) and the Warcup Fungal ITS database (Deshpande et al. 2016; hereafter, WITS). Restricting these databases to reads that included the FITS metabarcode locus (Epp et al. 2012) reduced species representation in the UNITE and WITS databases by 35.1% and 26.1% respectively (Table S3.1). Results show *CRUX* FITS contains more than 7 times as many species as does the R-UNITE database, and more than 16 times as many as the R-WITS database (Table S3.1). To assess phylogenetic coverage of *CRUX* FITS and the R-UNITE and R-WITS databases, we mapped reference databases onto a recently published phylogeny of Fungi (Choi and Kim 2017; Appendix 3.4). Results showed that *CRUX* FITS contains 88% of fungal species in this phylogeny whereas R-UNITE and R-WITS capture between 51-58% of fungal species (Figure S3.2; Table S3.6).

In reciprocal database comparisons, nearly all reads in the R-UNITE and R-WITS databases align to reads in *CRUX*-FITS. While the majority of reads in *CRUX*-FITS align to reads in R-UNITE and R-WITS, there are >42,000 *CRUX*-FITS sequences (6 - 9% of reads) that do not align to sequences in R-UNITE or R-WITS databases (Table S3.3). To determine the origin of these unaligned sequences, we again used HMMER (Finn et al. 2011) to build a profile hidden

Markov model of FITS sequences from BOLD (123,247 ITS sequences downloaded October 10, 2018).

We found that < 0.1% of unaligned reads in *CRUX*-FITS had significant hits to the hidden Markov model. However, BLAST searches indicate >98% of unaligned reads are indeed from the FITS metabarcode, suggesting that the majority of *CRUX*-FITS sequences match the metabarcode locus (Appendix 3.5.2). In contrast, only 89 reads in the R-UNITE and R-WITS databases did not align to *CRUX*-FITS (Table S3.3). BLAST searches of these reads indicate that all align to FITS genes. The TAXXI quality metrics indicate that *CRUX*-FITS yields much more reliable taxonomy at family, genus, and species ranks relative to published databases (average TPR using *CRUX*-FITS = 80.1; average TPR rate using restricted published databases = 43.4; Table S3.3, S3.5).

*3.2 Databases used for performance comparisons*

We compared *CRUX-*generated metabarcode specific databases (*16S* V4, *18S*, *18S* V4, *18S* V8-9, *12S*, *CO1*, and *FITS*; see Table S1.1) against several published databases that target the gene or internal transcribed spacer region associated with each metabarcode locus.

The *CRUX*-generated and published databases comparisons are as follows. The *CRUX*-generated *16S* was compared with the *SILVA* 128 99% clustered *16S* FASTA and the majority_taxonomy_7_levels taxonomy databases (Pruesse *et al.* 2007; Quast *et al.* 2013), Greengenes March 2011 release (DeSantis *et al.* 2006) and RDP Release 11 databases (Cole et al. 2013; downloaded from the Mothur website; https://www.mothur.org/wiki/RDP_reference_files). The *CRUX*-generated *18S* databases were also compared with the SILVA 128 99% clustered 18S FASTA and selected the majority_taxonomy_7_levels taxonomy databases. We compared the filtered *CRUX-*generated *12S* with the “MitoFish” Complete and Partial Mitochondria database downloaded July 15, 2018 (Sato et al. 2018). The filtered *CRUX*-generated *CO1* database was compared to the “Midori” (Midori-LONGEST_1.1 SPINGO_format database; Machida *et al.* 2017), “P-H” CO1 (Porter and Hajibabaei, 2018), and the “CO-ARBitrator” (Heller et al. 2018) databases. The filtered *CRUX-*generated FITS database was compared with the “UNITE” (UNITE Community 2017; Qiime release clustered at 99% including singletons) fungal database and the “WITS” Warcup Fungal ITS database (Deshpande et al 2016; downloaded from the RDP 11.3 release).

*3.3 Assessment of overlapping species between CRUX-generated and published libraries*

Published databases included sequences that were considerably longer than the metabarcode locus, or were missing the target locus entirely (Table S3.1; Figure S3.1). To compare database coverage at CO1 (Leray et al. 2013), 12S (Miya et al. 2015), and FITS (White et al. 1990; Epp et al. 2012) metabarcode loci, we considered only reads from the published databases which mapped to the loci of interest. This was accomplished by first generating EMBL *BLASTn* seeds for a locus using *CRUX* database generation parameters discussed in Appendix 1.1. Each respective seed library was then blasted against the relevant published databases using *CRUX* standard parameters to capture the reads corresponding to the target locus. All reads were de-replicated by sequence identifier and the largest instance of a read per accession was chosen. No read filtration was done on these reads. This reduced the size of published reference databases to only those reads that match target metabarcoding locus allowing for specific comparisons of coverage for a particular marker. Corresponding taxonomy files were then generated using *entrez_qiime* (Baker, 2016) or available taxonomy files were retained and in some cases modified for use with the *Anacapa classifier*. The resulting databases are referred to as “restricted” published databases. Bowtie2 libraries were generated for all restricted libraries for use with the *Anacapa classifier.*

The total number of unique species found in each database was determined using the run_namecount.py script developed for the *TAXXI* pipeline (Edgar 2018). We determined the lengths of each read in the full length published database, the restricted published database, and for each read in the Filtered *CRUX*-generated reference database (Table S3.1 & Figure S3.1). Due to taxonomic naming irregularities between the NCBI taxonomy and the taxonomy assigned to *16S* and *18S* published databases, we choose to only compare the counts of species and genera found in and unique to the *CRUX*-generated and the full length published database (Table S3.7).

*3.4 Assessment of database overlap in phylogenetic diversity*

We compared the phylogenetic breadth of *CRUX*-generated and restricted published reference databases by mapping taxa found in each database onto phylogenetic trees using the Interactive Tree of Life (itol.embl.de/). Given that metabarcode loci are designed to broadly capture diversity, we choose three phylogenetic trees that are representative of diverse clades captured by the following metabarcodes: *CO1, 12S* or *FITS*. *CO1* reads were mapped to a coleopteran phylogeny representing 367 coleopteran genera and 6 outgroups (McKenna et al. 2015), *12S* was mapped to a teleost phylogeny with 1410 teleost species and 6 outgroup species (Betancur-R et al. 2013), and *FITS* was mapped to a fungal phylogeny with 244 fungal species and 71 protist outgroups (Choi and Kim 2017).

To map database reads to corresponding phylogenetic trees, we determined taxon overlap between the phylogenetic tree taxa and the taxa present in *CRUX*-generated (*CO1, 12S*, or *FITS*) and the restricted published reference databases (R-Midori, R-CO-ARBitrator, R-P-H, R-MitoFish, R-UNITE or R-WITS). This was done by determining the NCBI taxon id for the genus or species in the database and matching it to the genus or species id of a phylogenetic tree taxon (Figure S3.2). The number of taxa from reference databases that overlap with the phylogeny is reported for *CO1* in Table S3.2, *12S* in Table S3.4, and *FITS* in Table S3.6. We then mapped taxa from each database that overlap taxa in the corresponding phylogenetic tree using the Interactive Tree of Life (itol.embl.de/; Figure S3.2).

3*.5 Reciprocal database testing*

To determine differences in taxonomic assignment between filtered *CRUX*-generated and restricted published databases, we ran reciprocal database tests between pairs of *CRUX*-generated and restricted published references databases for the following metabarcode loci: *CO1, 12S,* and *FITS*. For a given database pair, one database was used as the “query” database to be classified using the other “subject” database and vice versa. For example, *CRUX*-generated *CO1* was first used as the query database and was classified by the R-Midori subject database. Then R-Midori was used as the query database and was classified by the *CRUX-*generated *CO1* subject database.

*3.5.1 Using the Anacapa classifier to determine Bowtie2 alignment and BLCA taxonomic classification*

Using the *Anacapa classifier*, the query database was aligned to the subject database using *Bowtie2* (Table S3.3). As detailed in Appendix 1.3, query reads are first aligned to the subject database using Bowtie2 in end-to-end mode. The unaligned reads were then aligned using *Bowtie2* in local mode. All *Bowtie2* parameters are detailed Appendix 1.3. To reduce computation time, we retained only the best 50 returns rather than the best 100 returns. The *Bowtie2* returns were then run through the *Anacapa Bowtie2-BLCA classifier* using the default parameters with the following exceptions: percent of mismatch allowed between the query and subject for BLCA was set to (-b) 85 and the minimum percent of length of the subject relative to the query for BLCA was set to (-p) 90.

*3.5.2 Identification of unaligned database reads using BLAST*

We then tested all database reads that were unaligned during *Bowtie2* alignment to determine if they matched the target metabarcode locus (Table S3.3). For reciprocal test databases with fewer than 200 unaligned reads, for the *FITS* locus, and for all unmapped R-CO-Arbitrator reads and *CRUX*-generated *12S* reads that were not identified as belonging to the metabarcode locus using the hidden Markov models (HMM) described in section 3.5.3, sequence identity was verified with *BLASTn* searches. Only *BLASTn* hits with an e-value of 1e-06 of less were considered. If, for a given unmapped sequence, 5 or more of the top 10 hits matched the expected metabarcode locus, the sequence was considered positive for the locus. Matches to the locus were determined through parsing the subject titles in the *BLAST* output and looking for string matches to the metabarcode locus name. For example, for the FITS locus reads with the following names were accepted for the fungal ITS locus: “internal transcribed spacer” or “ITS”. For all other unaligned datasets, the locus was verified using HMM as indicated below in Appendix 3.5.3. HMM were not used for FITS as a suitable reference dataset could not be determined.

*3.5.3 Identification of unaligned database reads using profile hidden Markov models*

To identify the origin of *CRUX-12S* and *CO1* sequences that failed to map to the other reference databases, profile hidden Markov models (HMM) were used to search for homology between the putative metabarcode sequences and known annotated reference sequences. HMMs incorporate position-specific information on differential base conservation in DNA sequences, a characteristic that pairwise alignment methods such as *Bowtie2* and *BLAST* fail to take into account, and thereby provide added sensitivity in searching for remote homologues (*Eddy, 1998; Lagesen, 2007*).

For assessment of the unaligned 12S sequences from *CRUX*, barrnap version 0.9 (<http://www.vicbioinformatics.com/software.barrnap.shtml>) was employed to compare the putative *12S* sequences to the included HMM built from full-length aligned *12S* sequences from RefSeq. An e-value cutoff of 1e-06 was employed and hits with an alignment length less than 10% of the length of the HMM (corresponding to roughly 60% of the MiFish *12S* amplicon region) were rejected.

To assess the unaligned *CO1* sequences from reciprocal database testing, HMMs were again employed to compare the putative sequences to known reference sequences. *CO1* sequences from BOLD release 6.50 (http://v3.boldsystems.org/index.php/datarelease) were downloaded and MAFFT (Katoh et al. 2013) was used to perform a multiple sequence alignment (MSA) of the 163,325 reference *CO1* sequences. Subsequently, HMMER (*Finn et al. 2011*) was used to construct a profile HMM from the MSA using the *hmmbuild* command, and the HMM was searched against the unmapped *CO1* sequences from both *CRUX* and R-CO-ARBitrator using the *nhmmer* command. All of the unmapped *CO1* sequences were combined into a single fasta file to ensure that the e-values were directly comparable. An e-value threshold of 1e-06 was used to determine significant hits. Results of this analysis are shown in Table S3.3 and Figure S3.3.

3*.5.4 Assessment of classification using TAXXI metrics*

The *Anacapa classifier* taxonomic assignment results were assessed using the Taxonomic cross-validation by identity (TAXXI) framework of Edgar, 2018 (but also see *Edgar 2016* for more information on classification metrics*)*. Using scripts available from <https://drive5.com/taxxi/doc/index.html> and additional scripts written to run TAXXI with the *Anacapa classifier* (https://github.com/limey-bean/Anacapa), we determined the following classification metrics for each taxonomic rank. The true positive rate (TPR) which indicates how frequently the correct taxonomy was assigned out of the total number of opportunities for correct classification. The over-classification rate (OCR) which indicates how frequently too many ranks are predicted out of the total opportunities to make an over classification error. The under classification rate (UCR) which indicates how frequently too few ranks are predicted out of the total number of opportunities to make this error. The misclassification rate (MCR) which indicates how frequently a known name is incorrectly predicted out to the number of opportunities to make this error. The accuracy (ACC) indicates the number of correct taxonomic calls out of the number of opportunities to determine correct taxonomy. Note that only reads that align in *Bowtie2* are considered for TAXXI metrics. Results are reported for a *Anacapa Bowtie2-BLCA* bootstrap confidence cutoff of 60 (Tables S3.3, S3.5).

*3.6 Classification test to evaluate CRUX generated versus published databases performance on assignment of Sanger sequenced isolates*

We ran an additional benchmarking test of reference databases using Sanger sequenced isolates. We demonstrated the performance of *CRUX-*generated databases relative to published reference databases by assigning taxonomy with the *Anacapa classifier* to four Sanger sequenced isolate datasets (Table 2.2); 35 isolates sequenced for the *16S* v4 metabarcode locus (Callahan et al. 2016), 34 isolates sequenced for the *CO1* locus (Leray et al. 2017), four isolate pools of six - twelve species for the *18S* V4 locus and four isolate pools of six - twelve species for the *18S* V8-9 locus (Bradley et al. 2016). We classified these isolate datasets with databases for the appropriate locus (Appendix 3.2): five *16S* databases (*CRUX*-generated filtered and unfiltered, *SILVA* 128, Greengenes, and RDP), five *18S* databases (*CRUX*-generated filtered and unfiltered *18S* V4, *CRUX*-generated filtered and unfiltered *18S* V8-9, *SILVA* 128), and four *CO1* databases (*CRUX*-generated filtered and unfiltered *CO1*, R-Midori and R-P-H).

Results were summarized for five bootstrapping confidence cutoffs (60-95%). We then compared the total number of correct calls at levels of phylum through species and tallied the number of incorrect calls or empty classifications (i.e. where no assignment could be made). Z-tests in R (Team, 2000) were used to examine differences in the number of correct taxon paths reported at a bootstrapping confidence cutoffs of 60 across reference libraries*.* We used Bonferroni corrections to an initial alpha of 0.05 assess significance.

*3.6.1 Classification tests of Sanger sequenced isolates: filtered vs unfiltered CRUX-generated reference databases*

We found that the *CRUX* filtered databases lead to more correct taxonomic assignments than the other databases, with the exception of *CO1* unfiltered *CRUX* generated database (Figure S3.4 & Tables S3.8-11). The greater performance of the *CRUX* unfiltered *CO1* database was due to that three isolates were Platyhelminthes and the taxonomic assignment for Platyhelminthes was unresolved at higher taxonomy in NCBI, so references were not in the filtered database because they contained four consecutive NAs (e.g. Platyhelminthes;NA;NA;NA;NA;Platyhelminthes sp. FLTWO130-14 ).

*3.6.2 Classification tests of Sanger sequenced isolates: 16S comparisons*

For *16S*, the *CRUX*-generated *16S* filtered database identified the most correct taxonomic paths (Figure S3.4 & Table S3.8) at all bootstrapping confidence cutoffs (BCC 60-95; correctly identified 81.56 - 90.89%), relative to *SILVA128* (50.88 - 52.66%, Z-score = 12.18, p < 0.001), *Greengenes* (12.44 - 48.89%, Z-score = 13.73, p < 0.001), and the unfiltered *CRUX* database (1.56 - 22.89%, Z-score = 20.60, p < 0.001). The *CRUX* filtered library had the fewest erroneous calls identified for ranks in the taxonomic paths at all cutoffs (0 - 0.67%), relative to *SILVA128* 99% (30 - 36.4%), Greengenes (4.67 - 9.11%), and the unfiltered *CRUX* (0 - 7.33%).

*3.6.3 Classification tests of Sanger sequenced isolates: CO1 comparisons*

The *CRUX* generated *CO1* unfiltered database identified the most correct taxonomic paths (Figure S3.4 & Table S3.9) at all cutoffs (84.80 - 92.12%), relative to R-Midori (57.84 - 62.75, Z-score = 7.11, p < 0.001), R-PH (70.09 - 71.07%, Z-score = 5.49, p < 0.001), but not significantly greater than the *CRUX* filtered database (81.86 - 89.22%, Z-score = 1.02, p = ns). The *CRUX* unfiltered database identified the fewest erroneous calls to ranks in the taxonomic path (1.47 - 1.96%), followed by the *CRUX* filtered (3.92 - 4.41%), the R-Midori reference database (3.43 - 10.78%) and the R-PH reference database (24.5 - 26.9%).

*3.6.4 Classification tests of Sanger sequenced isolates: 18S comparisons*

*CRUX*-generated *18S* filtered database identified the most correct taxonomic paths (Figure S3.4 & Tables S3.10, S3.11) at all cutoffs (V4 50 - 87.5%; V8-9 56.94 - 86.11%), relative to *SILVA* (V4 19.4 - 19.4% Z-score 6.0374, p < 0.001; V8-9 18.04 - 22.22%, Z-score = 5.6864, p = 0.001), and the unfiltered *CRUX* database (V4 30.56 - 73.61%, Z-score 2.1056, p = ns; V8-9 31.94 - 68.05%, Z-score = 2.8062, p = 0.002). *CRUX* filtered database also had the fewer erroneous calls (V4 0 - 1.39%; V8-9 0 - 1.39%) compared to *SILVA* (V4 70.83 - 79.16 %; V8-9 50 - 65.27%, ) and the unfiltered *CRUX* (V4 0 - 6.94%; V8-9 0 - 6.94%).

*Figures*

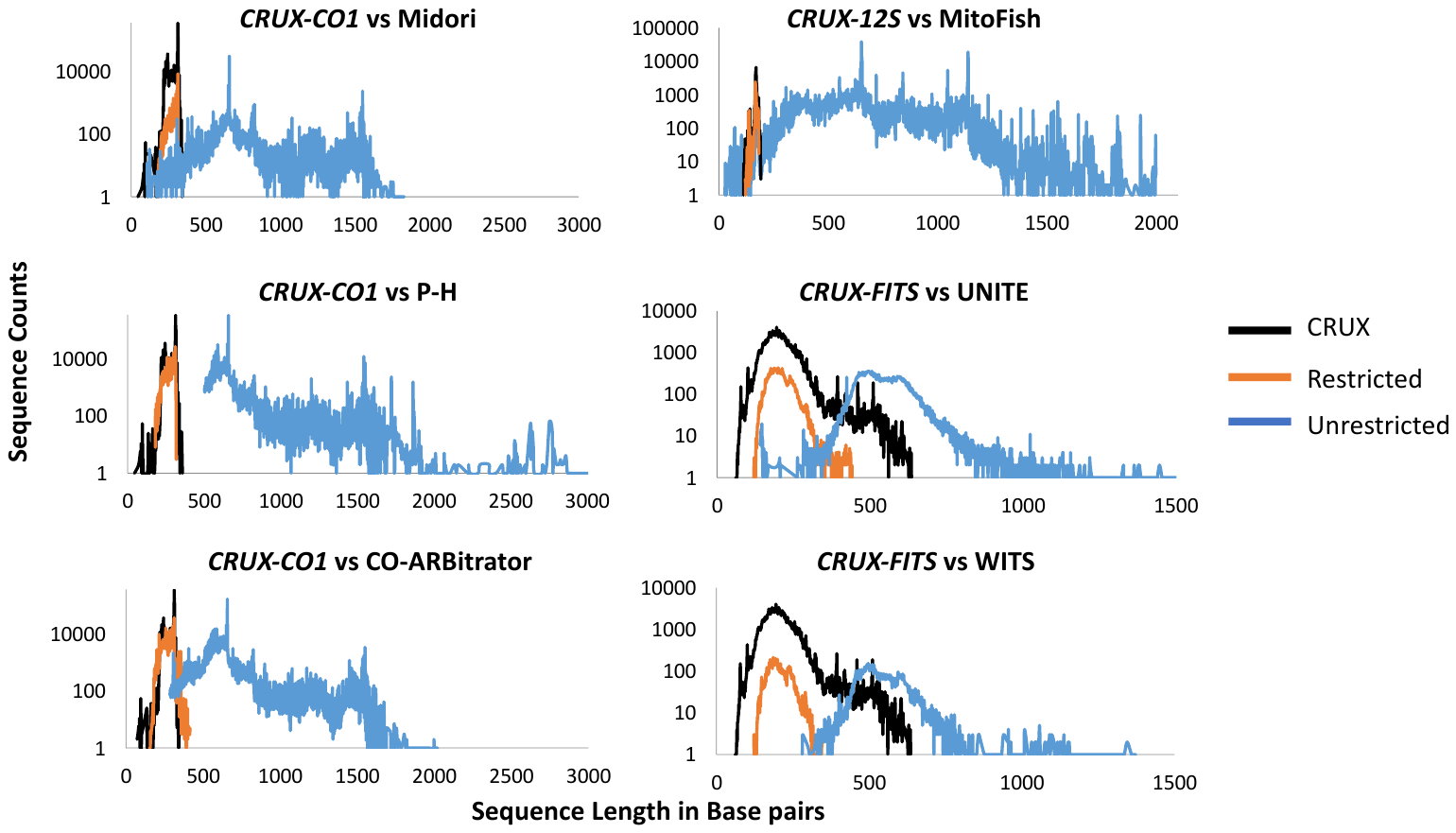

Figure S3.1. Length distributions of sequence reads in CRUX-generated and restricted and unrestricted published databases.

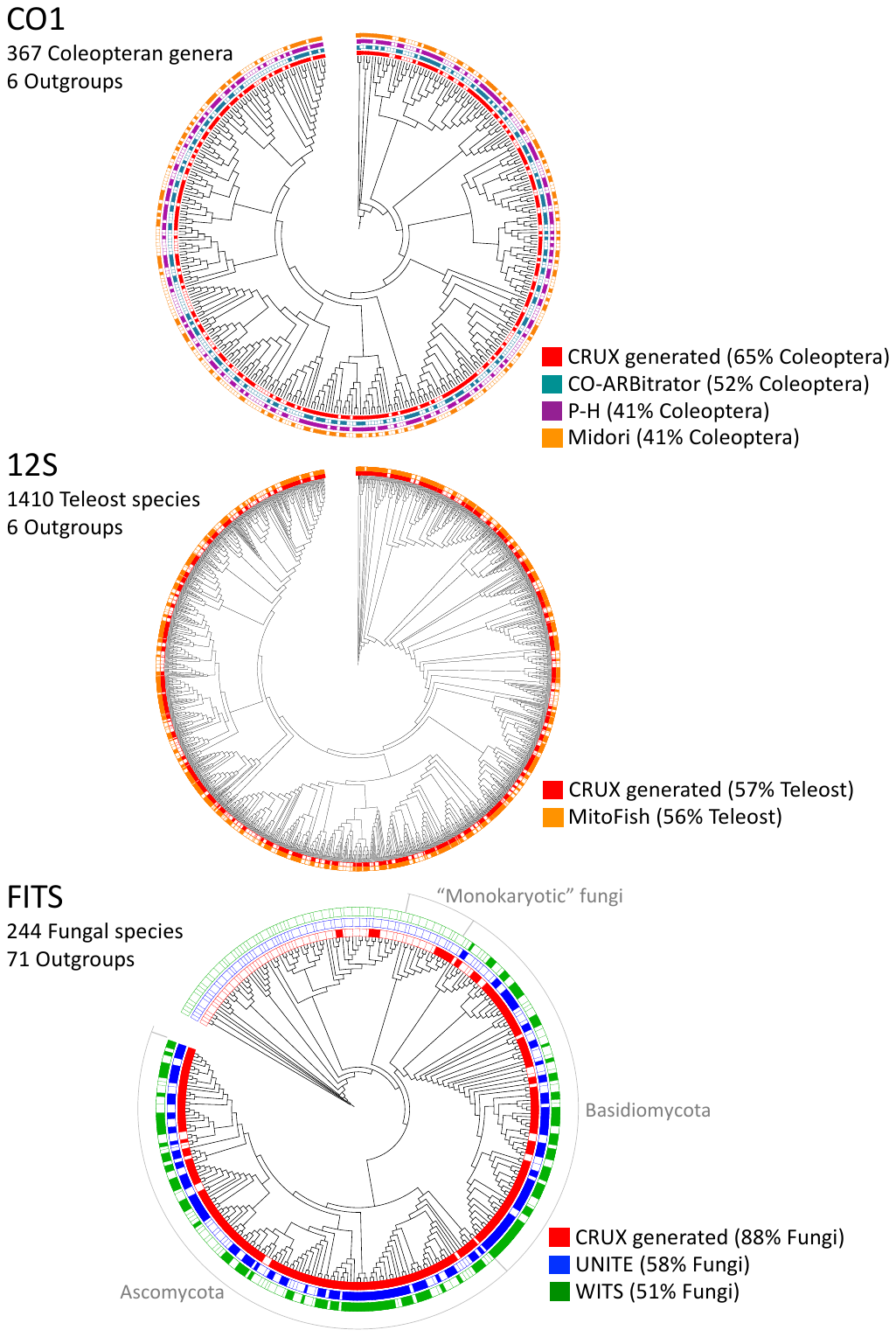

Figure S3.2. Comparisons of the phylogenetic breadth of CRUX-generated and restricted published reference databases. Indicated are the phylogenetic breadth of the Leray et al. (2013) CO1 barcode databases across a coleoptera phylogeny, the Miya et al. (2015) 12S barcode databases across a teleost phylogeny, and the Epp et al. (2012) FITS barcode databases across a Fungal phylogeny.

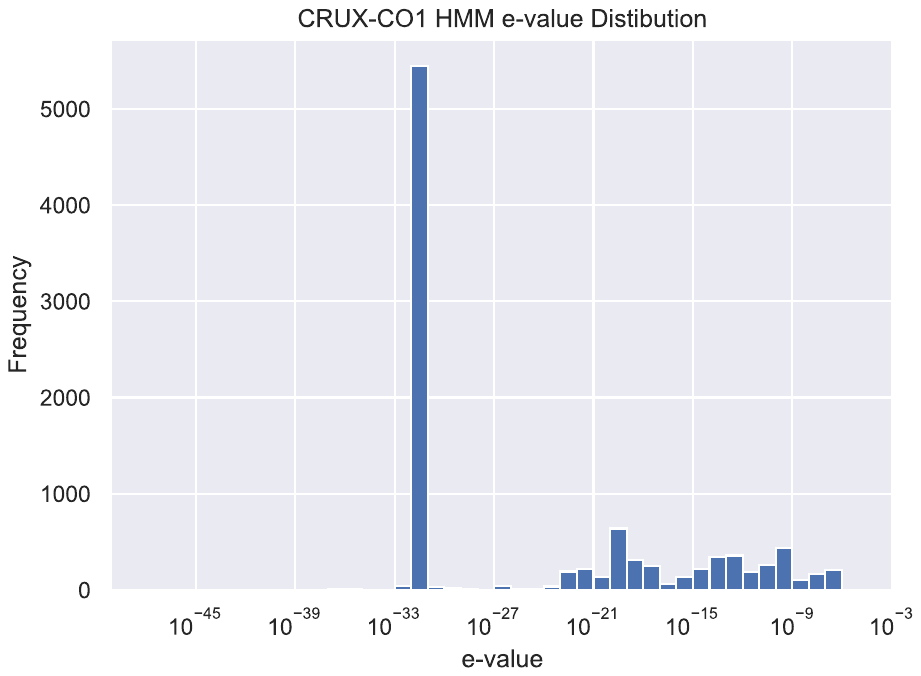

Figure S3.3. HMM e-value distribution of probable *CO1* reads for *CRUX-CO1* database sequences that did not align to R-Midori, R-P-H, or R-CO-ARBitrator databases.

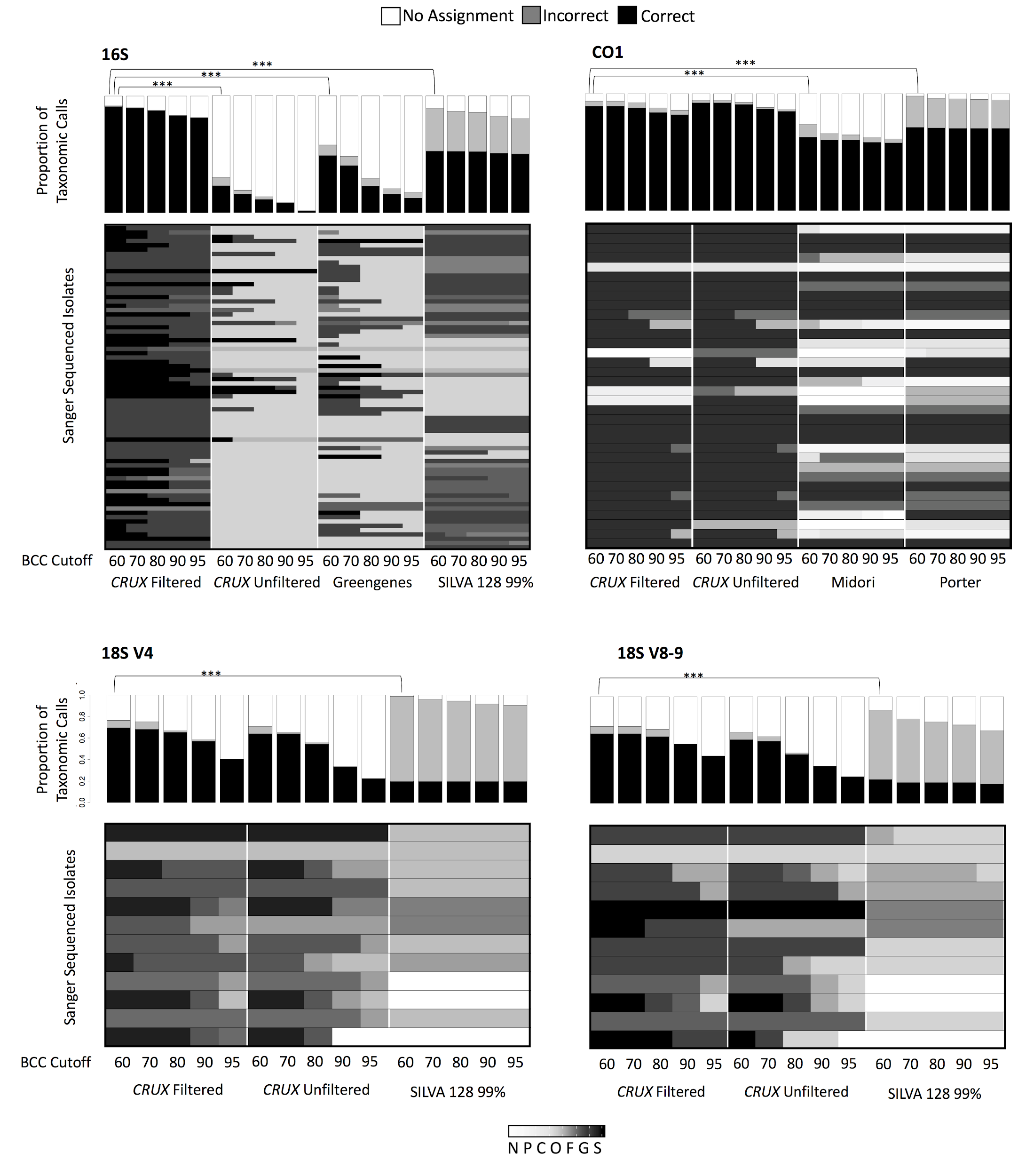
 Figure S3.4. Database comparisons between *CRUX* filtered and published reference databases for the *16S, CO1, 18S* V4 and *18S* V8-9 metabarcode locus regions. *16S* Sanger sequenced isolates (Krohn et al. 2016; Callahan et al. 2016; and Kozich et al. 2013; Appendix 2 Table 2) were assigned taxonomy using *CRUX* filtered and unfiltered, Greengenes, and the *SILVA* 128 99% percent clustered reference databases (Appendix 3.1). Isolates sanger sequenced for *CO1* (Leray et al. 2017; Appendix 2 Table 2) were assigned taxonomy using the *CRUX* filtered and unfiltered, the R-Midori “Midori”, and the R-P-H “Porter” *CO1* reference databases (Appendix 3.1). Sanger sequenced isolates (Bradley et al. 2016; Appendix 2 Table 2) for *18S* V4 and V8-9 were assigned taxonomy using the *CRUX* filtered and unfiltered, and the *SILVA* 99 percent clustered reference database (Appendix 3.1). The bar charts indicate the proportion of ranks in the taxonomic path were identified correctly, incorrectly, or not at all. The heat maps indicate the best correct taxonomic rank assigned to a given isolate for each database. The Best taxonomic rank for each database is given for bootstrap confidence cutoffs of 60 - 95%. The grey scale indicates the correctly assigned taxonomic rank: not correct (N) or assigned correctly to phylum (P), class (C), order (O), family (F), genus (G), or species (S). Z-scores between BCC 60 cutoff results are reported to a Bonferroni significance of ***=<0.001.
