## Supplementary material for "*Anacapa Toolkit*: an environmental DNA toolkit for processing multilocus metabarcode datasets"

***Appendix 4. Benchmarking the Anacapa Classifier module***

*4.1 Benchmarking the Anacapa Classifier module*

To verify that modifying the BLAST-*BLCA* classifier to accept *Bowtie2* alignments as input files (Appendix 1.3) does not change the reliability of taxonomic assignments, we used the Cross-Validation by Identity (CVI) approach of Edgar (2018) to compare the quality of taxonomic calls made using the original implementation of the BLAST-*BLCA* and the modified version in the Anacapa Classifier module. This approach explicitly models the variation between query sequences and the closest reference database entry (see Appendix 4.3). We used the TAXXI test and training sets, and Python scripts (https://drive5.com/taxxi/l) modified to run the Anacapa Classifier module. Anacapa’s *Bowtie2-BLCA* and BLAST-*BLCA* have highly similar CVI metrics to those reported by Edgar (2018). We also found that the Anacapa Classifier was generally faster at taxonomic assignment than the original implementation (Appendix 4.2; Table S4.1 & Figure S4.1), and that Anacapa’s *Bowtie2-BLCA* generally outperformed many commonly-used taxonomic classifiers using the TAXXI test-training datasets (Appendix 4.3; Tables S4.2-4).

Results of the CVI analysis showed that bootstrap confidence cutoffs can strongly influence quality of taxonomic calls generated by the Anacapa Classifier, and that optimal choice of cutoff may vary across metabarcodes (Tables S.2-4). For example, a bootstrap confidence cutoff of 80 had the highest accuracy for the 16S V3-V5 locus, whereas a cutoff of 60 optimized accuracy for the 16S V4 and WITS loci (Table S4.3). Thus, it is important to explore taxonomy tables generated by the Anacapa Classifier across a wide range of bootstrap confidence cutoffs. Although we do not present results on the sensitivity of taxonomy assignments to other parameters, the settings of “percent mismatch allowed between query and reference sequences” and “ minimum percent subject length relative to the query” (Appendix 1.3) likely also affect the quality of taxonomic assignments. As such, users should test their classifiers on training sets relevant to their study system to explore a range of parameters (see scripts designed for this purpose on the *CRUX* GitHub )

*4.2 Time benchmarking for the Anacapa Bowtie2-BLCA classifier relative to the BLAST-BLCA classifier*

To compare sequence handling time between the *Anacapa Bowtie2-BLCA* and *BLAST-BLCA* (Gao et al. 2017) classification methods, we conducted time benchmarking tests with sequence reads from field-collected *16S* and *CO1* datasets from Laguna (MiSeq data) and Stunt Ranch Santa Monica Mountains Reserve (HiSeq data; Table S2.3). We removed singleton ASVs, and for both methods, generated taxonomic assignments with the same parameters: 100 top returns, (-b) 80% percent length of subject relative to query, (-p) 80% percent of mismatch between query and subject, and 100 boot strapped replicates. For *Anacapa*, only “End-to-End” alignments were used. We tested unpaired forward, unpaired reverse, and merged ASVs, but not unmerged ASVs because they could not be assigned taxonomy with *BLAST-BLCA*. Classification assignments were based on filtered *16S and* *CO1* *CRUX*-generated reference databases. All *CRUX* generated reference databases were converted into BLAST libraries and taxonomy files were formatted for compatibility with *BLAST-BLCA*. Time for taxonomic assignment was recorded. Since each method processes unassigned reads differently, if a read was unassigned in either method, that read was removed from the analysis. We plotted the time to classify ASVs using for each method using scatter plot with R ggplot (Wickham 2016; Figure S4.1) and calculated regression summary statistics (see Table 4.1).

*Bowtie2-BLCA* performed significantly faster (Table S4.1 & Figure S4.1) than *BLAST-BLCA* (p<0.001 for all comparisons except forward only MiSeq Read)*. Anacapa’s* *Bowtie2-BLCA* classifier was particularly fast at assigning taxonomy when few best hits were returned or when the reference database was large. For example, the Anacapa *Bowtie2-BLCA* classifier had a shorter handling time for CO1 reads as compared to 16S reads and the CO1 database is 2.18x larger than 16S. We also found that with the Anacapa *Bowtie2-BLCA* classifier processed shorter HiSeq read lengths faster that longer MiSeq reads for both markers. In summary, both BLCA methods can perform rapid assignment.

*4.3 Comparisons of the Anacapa Bowtie2-BLCA and BLAST-BLCA Classifiers using Cross Validation by Identity*

To determine the performance of the *Anacapa Bowtie2-BLCA* and the BLAST-*BLCA* algorithm we followed the Cross Validation by Identity (CVI) methodology implemented in TAXXI (Edgar 2018; <https://drive5.com/taxxi/doc/index.html>). Edgar (2018) used this method to evaluate taxonomic classification for 20 methods including *BLAST-BLCA*. This involved running classifier methods on test datasets using training datasets derived from full length *16S*, *16S* V4 and *16S* V3-5 regions, and full length Fungal ITS (<https://drive5.com/taxxi/doc/fasta_index.html>) sequences. Edgar (2018) modified each dataset to standardize the number of reads included per genera, which reduced computational requirements. In addition, datasets were clustered to different percent identities (100, 99, 97, 95, and 90) to simulate a classifiers ability to detect novel taxa. Metrics used to evaluate the performance of each classifier are described above in Appendix 3.4.4*,* and include over classification rate (OCR), under classification rate (UCR), misclassification rate (MCR), true positive rate (TPR), and Accuracy (ACC).

We used *TAXXI* scripts and custom scripts to compare *Bowtie2-BLCA* and BLAST-*BLCA* on UCLA’s Hoffman2 HPC. Scripts can be found at <https://github.com/limey-bean/Anacapa>. Both *BLCA* classifiers were run using the default parameters of BLAST-BLCA (https://github.com/qunfengdong/BLCA), which were also used in Edgar 2018: BLCA considered the top 50 alignments, 90% percent of mismatch allowed between the query and subject, 85% percent length of subject relative to query, and 100 bootstrap replicates. We compared the two classifiers at bootstrap confidence cutoff of 80% used by Edgar (2018) as well as 100, 90, 70, 60, 50, and 40. Both classifiers were run 3 times per database and we reported the average and standard deviation (Tables S4.2-4). We also include taxonomic assignment metrics for other classifiers tested in the Edgar 2018 paper in Tables S4.3, S4.4.

*Figures*

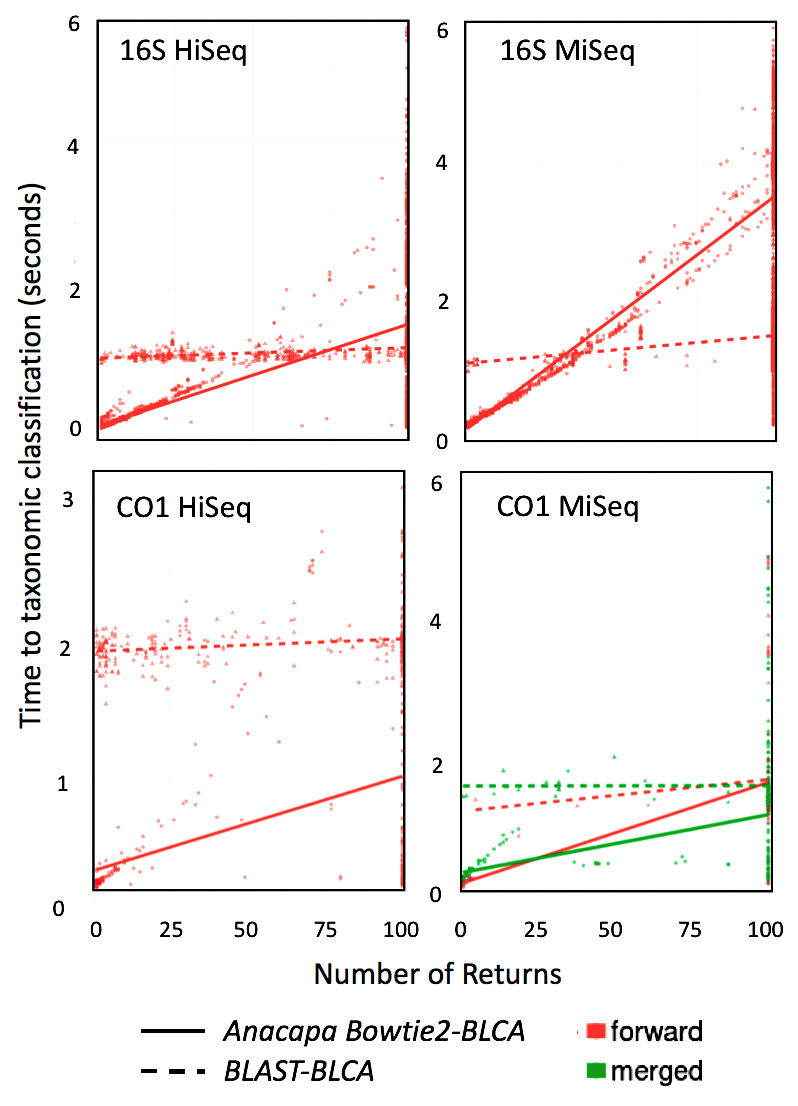

Figure S4.1. Differences in sequence handling times taxonomic classification of ASV using *BLAST-BLCA* and *Anacapa Bowtie2-BLCA classifier* methods. The number of returns indicates the number of subject alignments to the each query Amplicon Sequence Variant (ASV) generated by the *Anacapa Bowtie2-BLCA* or *BLAST-BLCA* classifiers for *CO1* or *16S* metabarcode loci*.* Sequence data is from three Laguna (MiSeq) and three University of California Stunt Ranch Santa Monica Mountains Reserve (HiSeq) libraries (Appendix 2 Table 3). Only non-singleton ASVs were used, and only ASVs in which a taxonomic assignment was given for both methods at a bootstrap confidence cutoff of 60% or greater. This reduced the dataset to primarily forward reads because few reverse and merged reads passed these criteria. Summary statistics are in Appendix 4 Table 1.
