## Supplementary material for "*Anacapa Toolkit*: an environmental DNA toolkit for processing multilocus metabarcode datasets"

***Appendix 5. Processing MiSeq and pseudo-HiSeq data with the Anacapa Classifier***

*5.1 Anacapa’s ability to Process MiSeq and pseudo-HiSeq data*

To assess the effect of read length on the ability of the *Anacapa Toolkit* to assign taxonomy in eDNA studies, we trimmed 250 or 300 bp (MiSeq-length) reads in silico to 150 bp (HiSeq-length) fragments, effectively creating matching “pseudo-HiSeq” datasets (Appendix 5.2). For this test, we selected published MiSeq datasets generated by characterizing known communities for CO1 (n=1), 16S (n=1), or 18S-V4 (n=4) / V8-9 (n=4) metabarcodes (Appendix 2.3). We used cutadapt to generate pseudo-HiSeq version of each of these datasets by trimming sequences to the first 150 bp. We processed both untrimmed and trimmed datasets using Anacapa’s Quality Control and Classifier modules and used *CRUX*-generated reference databases for classification. We then performed linear regressions to compare read counts assigned to each taxon in MiSeq-length and pseudo-HiSeq datasets (Appendix 5.3). The *Anacapa Toolkit* can consistently assign taxonomy to MiSeq-length and HiSeq length reads, except in cases where 150 bp reads are too short to cover diagnostic sequences. For 16S-V4, the number of reads assigned to each taxonomic call in pseudo-HiSeq and MiSeq length datasets was highly correlated when taxonomy was considered at the species level (R2 = 0.58), 327 and identical (R2=1) at the genus level (Figure S5.1; Table S5.1, S5.2). In seven of the eight 18S isolate pools used in this comparison (Appendix 5 Tables S5.3,S5.4), Anacapa generated similar taxonomic calls at the species level for MiSeq and pseudo-HiSeq datasets (R2> 0.65).

The *Anacapa Toolkit* assigned less well-resolved taxonomy for many sequences in the pseudo-HiSeq version of one of the 18S isolate pools (18S V8-V9 M61, Figure S5.1) because diagnostic barcode sequences fell outside of the first 150 base pairs. Finally, the *Anacapa Toolkit* assigned similar taxonomy to MiSeq and pseudo-HiSeq versions of the CO1 datasets, except for one taxon, which was assigned to the rank of species with MiSeq sequences and only to the rank of class with HiSeq-length sequences (Figure S5.1 & Table S5.5). Together, these results demonstrate that the *Anacapa Toolkit* can process MiSeq and HiSeq reads, and that the depth of taxonomic resolution is dependent on target barcode length and taxa of interest.

*5.2 Anacapa functionalities with MiSeq and pseudo-HiSeq data*

*5.2.1 Mock datasets*

The *Anacapa toolkit* was designed to assign taxonomy to paired-end sequence reads that are long enough to merge, but also to those reads that are not long enough to merge. To demonstrate that unmerged paired reads contain useful and comparable data to merged paired-end reads, we generated pseudo-HiSeq data sets from sequenced isolate pool data generated on a MiSeq (2 x 250 or 2 x 300 bp read lengths; Table S2.2). These datasets were truncated to the maximum read length generated by an Illumina HiSeq (2 x 150 bp) by trimming the 3’ read ends using *cutadapt* (Matrin, 2011). The majority of the pseudo-HiSeq reads were not long enough to merge (Appendix 5 Tables 5.2-5).

*5.2.2 Processing the mock data*

The MiSeq and pseudo-HiSeq data sets were run through the *Anacapa Sequence QC and ASV Parsing* and the *Anacapa BLCA classifier* modules using the methods described in Appendices 1.2 and 1.3. All singleton ASVs (ASVs that represent a single read in the entire dataset) were removed from subsequent analyses.

*5.3 Anacapa Analysis of MiSeq and pseudo-HiSeq data*

We used linear regression (R Core Team, 2017) to test the correlation in taxonomic assignments for MiSeq and pseudo-HiSeq data sets given a bootstrapping confidence cutoff of 60. We ran this analysis for read counts of the entire taxonomic path given for each dataset, and also for read counts the taxonomic paths to the rank of genus. We found moderate to strong correlations in taxonomic assignments between MiSeq and pseudo-HiSeq data for the mock community datasets (Figure S5.1; Table S5.1), we found Illumina read length affects the distribution of ASV types (paired merged, paired unmerged, forward, reverse) assigned for a data set (Tables S5.2-5), and the number and count of ASVs recovered for a given taxonomic path (Tables S5.6-9).

*Figures*

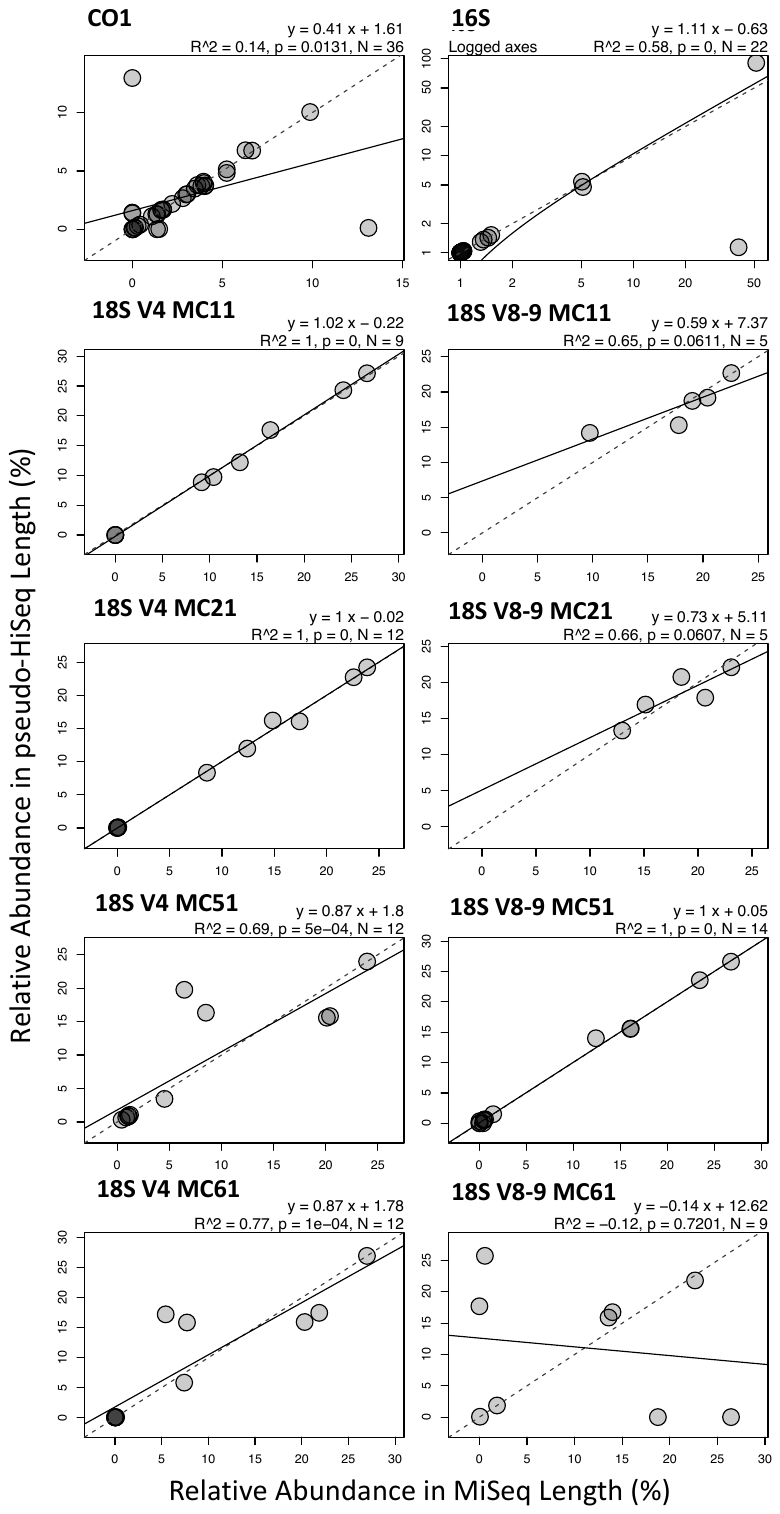

Figure S5.1. Expected 1:1 versus linear regressions of MiSeq vs pseudo-HiSeq read counts for genera in the mock datasets: 35 pooled isolates amplified for the *16S* metabarcode (Callahan et al. 2016; Appendix 2 Table 2; Appendix 5 Table 4 and 6), 34 pooled isolates amplified for the *CO1* metabarcode (Leray et al. 2017; Appendix 2 Table 2; Appendix 5 Table 3 and 7), and up to 12 pooled isolates amplified for the *18S* metabarcodes V4 and V8-9 regions (Bradley et al. 2016; Appendix 2 Table 2; Appendix 5 Tables 4-5 and 8-9). Each graph includes a dotted 1:1 line that would indicate a perfect correlation in taxonomic assignment between datasets. Overlapping circles appear darker. Note logged axes in the *16S* plot.
