## Supplementary material for "*Anacapa Toolkit*: an environmental DNA toolkit for processing multilocus metabarcode datasets"

***Appendix 6. Demonstration of Anacapa with CALeDNA field-collected samples***

*6.1 Generation of eDNA metabarcode sequence libraries from California samples*

30 samples collected for CALeDNA were used in this study (Table S6.3). These kelp forest water samples were collected as 1L seawater samples from 10m depth on SCUBA by the Channel Islands National Park Service (June, 2016) using sterile collapsible bags at three sites around Anacapa Island. Sea water was gravity filtered onto 0.2um Sterivex filters for 40 minutes. Sample were stored at -20°C for seven days prior to extraction.

*6.1.1 DNA extraction*

Genomic DNA from kelp forest samples was extracted from the filter with using the Qiagen DNeasy Blood and Tissue kit (Qiagen, Valencia, CA, USA) following the modifications of Spens et al. (2017). The DNA was stored at -20ºC.

*6.1.2 Sequence library generation*

The kelp forest eDNA was amplified by polymerase chain reaction (PCR), using the *12S* and *CO1* primers in Table S2.2 but with one modification. Primers were synthesized with the addition of a 5’ linker that matches the region of the Nextera transposome and matches Nextera Indices (Illumina, San Diego, CA, USA). PCR amplification was performed in triplicate using a 25 μL reaction mixture containing 12.5 μL of QIAGEN Multiplex Taq PCR 2x Master Mix, 6.5 μL of dH2O, 2.5 μL of each primer (2 μmol/L each), and 1 μL of template DNA.

The PCR cycling for the both amplicons was conducted in a thermocycler by a touchdown program: initial denaturation at 95°C for 15 min, 13 cycles of denaturation at 94°C for 30 sec, beginning annealing at 69.5°C for 30 sec (temperature was decreased by 1.5°C every cycle until 50°C was reached), and extension at 72°C for 1 min. 35 additional cycles were carried out at an annealing temperature of 50°C, followed by a final extension at 72°C for 10 min.

The PCR products were run on 2% agarose gels to ensure correct product size. The replicates were then pooled by sample. Within each sample, metabarcode amplicons were pooled by copy number. This was accomplished by cleaning amplicon DNA with SPRI beads (Applied Biological Materials, Vancouver, British Columbia, Canada) and the quantifying amplicon DNA with Quant-iT™ dsDNA Assay Kit, high sensitivity (Thermofisher Scientific, Waltham, MA, USA) on a Victor3 plate reader (Perkin Elmer Waltham, MA, USA). We pooled amplicons evenly by amplicon copy number calculated using the following formula: number of amplicon copies = ( concentration of DNA * 6.022x10^23^) / (length of the amplicon * 1x10^9^ * 650).

Sample DNA libraries were prepared using the Nextera Index A and or D Kit (Illumina, San Diego, CA, UCA) and KAPA HiFi HotStart Ready Mix (Kapa Biosystems, Wilmington, MA, USA). The PCR was performed using a 25 μL reaction mixture containing 12.5 μL of Kapa HiFi Hotstart Ready mix, 0.625 μL of primer i7, 0.625 μL of primer i5, and approximately 5ng of template. The PCR cycling for the indexing was conducted in a thermo-cycler: denaturation at 95˚C for 5 min, 5 cycles of denaturation at 98˚C for 20 sec, annealing at 56˚C for 30 sec, extension at 72˚C for 3 min, followed by a final extension at 72˚C for 5 min.

The indexed PCR products were run on 2% agarose gels to ensure correct product size. Libraries were then bead cleaned and DNA concentration was quantified as described above. Indexed libraries were pooled in equimolar concentration, and sent to the QB3-Berkeley FGL (University of California, Berkeley, CA, USA) where they were run with other samples on a MiSeq Reagent Kit V3. Target read depth per metabarcoding locus per sample was 100,000 reads. 30% PhiX was added to the sequencing runs.

*6.2 Real data sequence processing*

The datasets were run through the *Anacapa Sequence QC and ASV Parsing* and the *Anacapa classifier* modules using the default parameters. All singleton ASVs (ASVs that represent a single read) were removed from subsequent analyses. ASVs (unpaired forward, unpaired reverse, and merged) for *16S* and *CO1* samples were also assigned taxonomy with *SKLearn* as described in Appendix 4.

*6.3 Diversity Analysis*

Species and genus richness was calculated by summing the number of unique genera and species for each sample. Due to the presence of unassigned species and genera assignments (e.g. Family Diptera or Salispi sp.) species and genus richness counts were inflated by 1 (e.g. NA). Species and genus richness was compared between classifiers using ANOVA, implemented in R using the *vegan* package (Oksanen et al. 2013). We also calculated the Jaccard similarity distance for samples at both the species level and genus level grouping using the *phyloseq* package in R (McMurdie and Holmes, 2018). We compared community composition between classifiers and habitats using the PERMANOVA (Adonis) using *vegan*. We ran homogeneity of dispersion tests between classifiers and habitats the betadisp function from *vegan* in R.

*6.4 Results of California field-collected eDNA samples processed with Anacapa*

*6.4.1 Explanation of 12S taxonomy assignment*

A detailed taxonomic summary of the CALeDNA *12S* samples can be found in the demo dataset of the <https://gauravsk.shinyapps.io/ranacapa/> and in Table S6.1. The *ranacapa* demo dataset also includes *CO1* reads for these samples (Table S6.2). Many California vertebrates have not been sequenced for the *12S* locus. Many of the coastal marine species in this region have not been sequenced for the 12S locus, resulting in assignments to genus level only, or to non-Californian sister species For instance, an ASV assigned to *Sardinops melanostictus*, a fish native to Japan (www.fao.org/), was likely DNA from a native California Sardinops species, but only *Sardinops melanostictus* was in the CRUX-12S database. The Anacapa Toolkit successfully captured a broad diversity of fish taxa, including keystone kelp forest species like California sheephead, *Semicossyphus pulcher*. Despite being designed for fish, the 12S MiFish barcode also captured other vertebrate taxa like California sea lion, *Zalophus californianus* a species of management concern, and rare and transient species like the booby seabird, *Sula* sp. In several instances we found eDNA sequences that match the correct genus but not the correct species, due to the absence of *12S* sequence for that species in Genbank. Four of the genus-level identifications had only one species known to occur in California, so we annotated assignments as the California species. There were five cases in which we identified species not found in California, but that are known sister species of a Californian species. No published *12S* DNA barcodes were available for those Californian taxa and the reads that were assigned to sister taxa are most likely derived from the Californian taxa. The Marbled blenny *Paraclinus marmoratus* was the only species misidentified that had any matching partial reference sequence (NCBI Accession Version U90367.1). Six of the species identified matched to non-native Southern California fish species, but are know invasive species. Presence of human may be a result of detection of the SCUBA divers collecting the samples or other divers in the surrounding kelp forest.

*6.5 Figures*

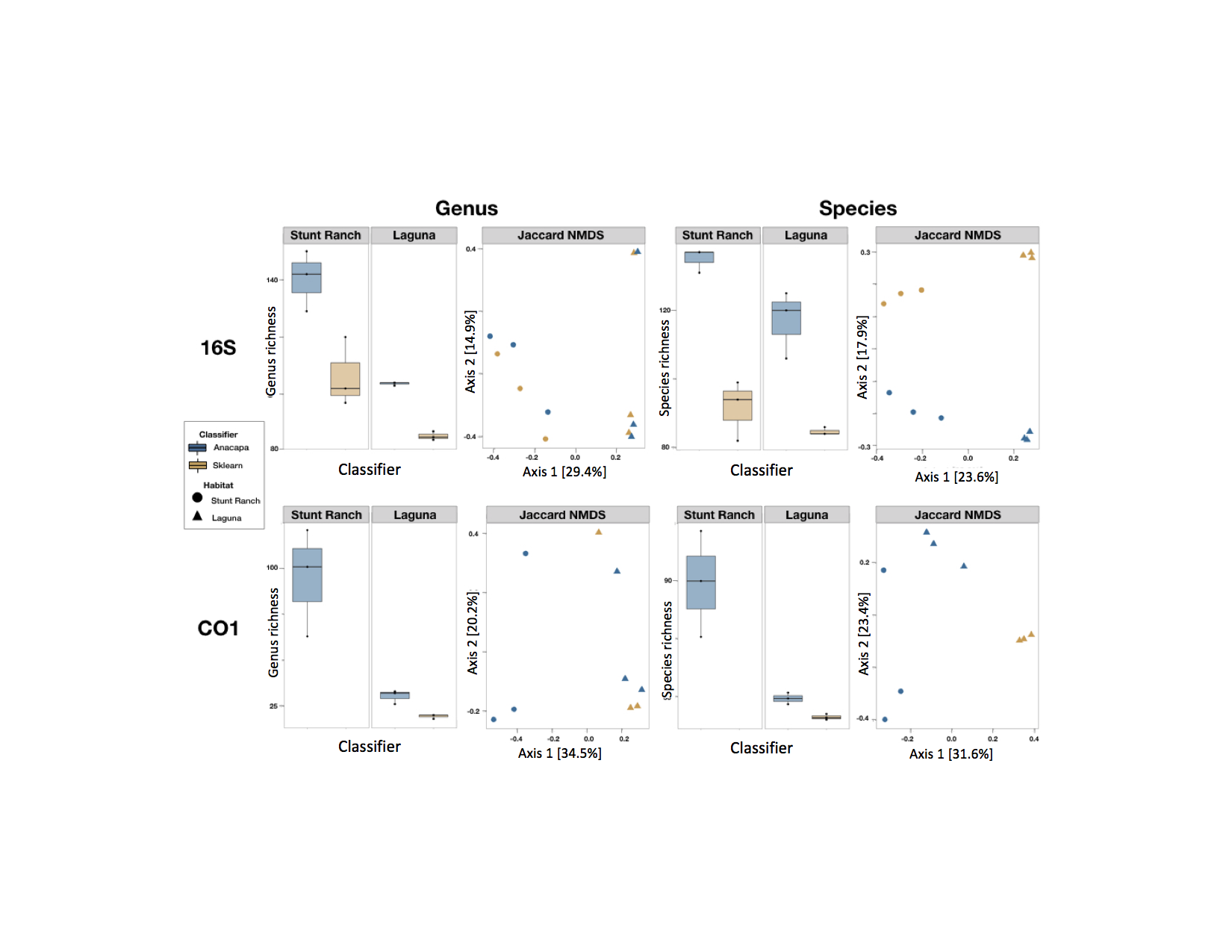

Appendix 6 Figure 1. Alpha and Beta Diversity Comparisons of *Bowtie2-BLCA* and *Qiime2* *SKLearn* classifiers on California field-collected eDNA samples. Here we compare the classifiers on three eDNA surface soil samples collected from University of California Stunt Ranch Santa Monica Mountains Reserve and the Laguna Coast Wilderness Park for two barcode loci: *16S* and *CO1*. We found that the *Anacapa Bowtie2-BLCA classifier* identifies significantly more genera and species level assignments than the *SKLearn classifier* for the *16S* marker (ANOVA, genus: p <0.034, species: p<0.009).
